# Early establishment and life course stability of sex biases in the human brain transcriptome

**DOI:** 10.1101/2024.10.11.617734

**Authors:** Clara Benoit-Pilven, Juho V. Asteljoki, Jaakko Leinonen, Juha Karjalainen, Mark J. Daly, Taru Tukiainen

## Abstract

To elaborate on the origins of the established male-female differences in several brain-related phenotypes, we assessed the patterns of transcriptomic sex biases in the developing and adult human forebrain. We find an abundance of sex differences in expression (sex-DE) in the prenatal brain, driven by both hormonal and sex-chromosomal factors, and considerable consistency in the sex effects between the developing and adult brain, with little sex-DE exclusive to the adult forebrain. Sex-DE was not enriched in genes associated with brain disorder, consistent with systematic differences in the characteristics of these genes (e.g. constraint). Yet, the genes with persistent sex-DE across lifespan were overrepresented in disease gene co-regulation networks, pointing to their potential to mediate sex biases in brain phenotypes. Altogether, our work highlights the prenatal development as a crucial timepoint for the establishment of brain sex differences.

## Introduction

A variety of human morphological, physiological, and behavioral phenotypes differ between males and females. For instance, sex differences in the structural and functional brain organization, e.g., volume and neuronal connectivity, have been uncovered^1–3^. Furthermore, numerous neurological and psychiatric disorders display well-established differences in incidence, prevalence, severity, and/or age at onset between sexes^4^. For example, autism spectrum disorder (ASD) shows clear sex biases in its prevalence (4.5:1 male-to-female ratio) and severity (more female ASD patients presenting with intellectual disability)^5,6^. Despite these abundant sex differences in phenotypes that relate to the function of the human brain, little is known about how and when they appear.

Although genetic factors often explain a large fraction of the phenotypic variability, there is limited evidence that genetic architecture has a substantial contribution to sex differences in complex traits^7,8^. For brain-related traits and diseases, genetic studies have found only minor differences both in common variant associations^9^ and in rare variant burden^10,11^ between males and females. For ASD, there is evidence for the difference in the liability threshold between sexes playing a part in the difference in prevalence, as females require a higher genetic load to meet the diagnostic criteria for the disease^12,13^. However, as genetic studies have generally been unsuccessful in explaining the abundance of phenotypic sex differences, numerous studies have searched for alternative explanations for the observed divergences between males and females at other levels of regulation, i.e., in epigenetics^14,15^ and transcriptomics^16^.

In contrast to the limited sex differences in the genetic regulation of phenotypes^8,16^, a growing number of transcriptomic studies point to the abundance of sex-differentially expressed (sex-DE) genes in the adult brain^16–20^. However, as gene expression throughout the lifespan is very dynamic^21^, these transcriptomic sex differences observed in the adult brain may be distinct from the processes leading to sex biases in phenotypes, as many neurodevelopmental disorders, including ASD and attention-deficit/hyperactivity disorder (ADHD), are rooted in fetal development^22,23^. Gene expression sex biases in the prenatal brain have also been described^17,18,24–27^, yet most studies investigating these have been limited by the small sample sizes and the heterogeneity of the data, including large variation of gestational ages and brain regions sampled. One of the largest studies thus far by O’Brien and colleagues^25^ analyzed 120 prenatal whole brain samples from the second trimester of pregnancy, discovering more than 2000 differentially expressed genes between males and females. Such a large number of sex-DE genes uncovered suggests sex biases in brain gene expression appear very early during fetal development. Further, it remains unclear if these differences are transient or persist into adulthood.

The brain tissue is very complex with specific regions displaying distinct characteristics, functions, and transcriptomic patterns of expression^28,29^. The forebrain is one of the three primary vesicles of the brain during development, and it can be identified as early as 3-4 weeks post-conception (PCW) in humans^30^. This vesicle gives rise to the cerebral cortex, which composes around two-thirds of the adult brain’s total mass, as well as to the amygdala, the hippocampus, the hypothalamus, and the basal ganglia. The forebrain is implicated in the higher brain functions, such as thinking, memory, and sensory processing (e.g., visual, auditory), and is affected by neuroanatomical alterations in several neurological diseases, including ASD^31,32^ or schizophrenia^33^. For all these reasons, the forebrain is a particularly interesting brain region to study sex differences.

In this study, we explored the extent, dynamics, and biology of transcriptomic sex differences in the developing and adult human forebrain to provide insights into the timing and mechanisms of the widespread brain-related male-female differences. To this end, we used RNA sequencing (RNA-seq) datasets from the early prenatal, i.e. first and second trimester, and adult forebrain totaling together 1899 samples. Through differential expression analyses comparing male and female samples, we assessed the patterns of sex-DE in and between prenatal and adult forebrain and then investigated the potential causes and consequences of these transcriptomic sex differences. We show that sex-DE genes 1) are abundant in the human forebrain, in particular in the prenatal brain, 2) are found already very early during brain development, and 3) are involved in several important developmental processes. Our analysis reveals a hormonal contribution to the prenatal-specific sex differences. We also demonstrate that, despite vast structural and molecular-level differences between the prenatal and adult brain, a substantial portion (30%) of the sex-DE genes is shared across both life stages, although the effect sizes were generally weaker in adulthood. These lifelong sex-DE genes include not only sex-chromosomal genes, including those that escape from X chromosome inactivation (XCI), but also autosomal genes related to neuronal functions. Overall, these observations highlight a substantial degree of functionally relevant male-female differences in the human forebrain. The emergence of these sex biases in early development points to biological underpinnings that can plausibly contribute to the sex differences detected in diverse brain-related phenotypes in humans.

## Results

### Description of datasets

To investigate the characteristics of sex biases in the human brain transcriptome, including temporal dynamics across distinct life stages, we analyzed the gene expression signatures in two forebrain datasets, one from the prenatal brain and the other from the adult brain. For the prenatal dataset, we extracted forebrain samples from the Human Developmental Biology Resource (HDBR) Expression dataset^34^ (see Methods) resulting in 266 RNA-seq samples (130 samples from 37 female individuals and 136 samples from 35 male individuals, with a median of 2 samples per individual) from the first and early second trimester (5-17 weeks post-conception (PCW)) pregnancies (**Figure 1A**; **Figure S1A**; **Table S1**). The adult brain samples were obtained from the Genotype-Tissue Expression project (GTEx v8)^35^. Again, we selected only samples coming from the forebrain region (see Methods), accounting for a total of 1633 RNA-seq samples (434 samples from 91 female individuals and 1199 samples from 246 male individuals, median of 5 samples per individual) (**Figure S1B**; **Table S1**). In addition to the forebrain region’s biological relevance to many neurodevelopmental diseases^36^, we focused on this brain region to ensure a dataset as homogeneous as possible (see Methods; **Figure S2A-C**). Our definition of males and females refers to genetic sex (i.e. XY and XX), determined via sex chromosome gene expression (see Methods).

**Figure 1:**
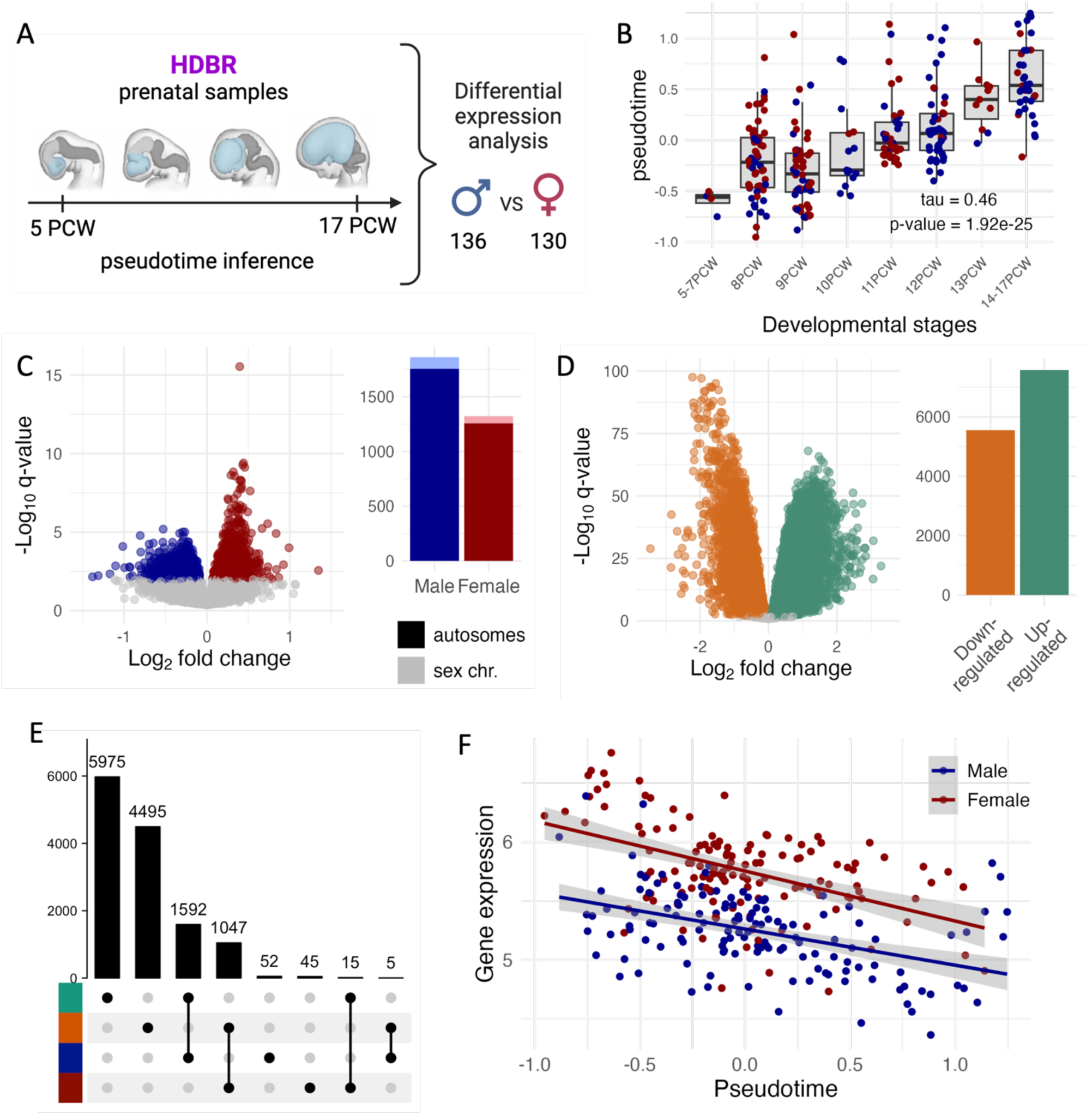
Differential expression (DE) analysis in the prenatal forebrain. (A) The prenatal data set consisted of 266 samples from the Human Developmental Biology Resource (HDBR) ranging from 5 to 17 post-conception weeks (PCW). We first carried out a pseudotime analysis to place the samples on a continuous developmental trajectory and then performed a differential expression analysis between the male and female samples to quantify sex-differential gene expression. (B) Correlation between the inferred pseudotime variable and the reported developmental stages in PCWs. Male samples are colored in blue, female samples in red. The Kendall’s tau rank correlation and its associated p-value are given in the plot. (C-D) Barplot of the number of DE genes (q-value<0.1) and volcanoplot for the two analyses: (C) sex-DE genes and (D) pseudotime-DE genes. On the sex-DE volcanoplot only autosomal genes are shown. Only significant genes are colored in both volcanoplots. (E) Intersection of the findings from the two DE analyses as an upset plot taking into account the direction of effect. The upper barplot shows the size of the intersections. (F) Expression of *ZFX* along pseudotime. *ZFX* is female-biased (q-value=4.5x10^-12^, logFC=0.52), and downregulated along pseudotime (q-value=2.4x10^-18^, logFC=-0.43), but does not show any significant interaction (q-value=0.42, logFC=0.02).

### Overview of the analyses

All analyzed prenatal and adult RNA-seq data was previously generated with Illumina technologies^34,35^. Gene quantification was carried out with Salmon^37^ for the prenatal samples, while we used the pre-computed gene counts for the adult data provided by the GTEx consortium (see Methods). 17614 and 17263 protein-coding and lincRNA genes were detected in the prenatal and adult data, respectively.

As prenatal development is a continuous process and the partitioning of the samples in discrete stages (i.e., post-conception weeks or developmental stages) is subject to variability, we modeled the developmental process as a continuous variable, which we will refer to as pseudotime (see next section), inferred from the gene expression measures. For methodological consistency, we carried out a similar pseudotime analysis in the adult dataset. Then, we conducted differential expression analyses comparing males and females in each data set separately using the estimated pseudotime as a covariate, followed by investigations into the (dis)similarity of the sex bias profiles in the pre- and postnatal brain. Finally, we assessed the biological relevance of these sex biases through several enrichment analyses, with the overall aim of providing a better understanding of the origins of brain-related sex differences.

### Pseudotime inference for prenatal forebrain samples

The prenatal samples from HDBR were collected from fetuses between 5 and 17 PCW from voluntary interruption of pregnancy. Although we focused our analysis on the forebrain, the age range of the samples (**Figure S1A**) covers a highly active developmental period^30^, resulting in transcriptional heterogeneity of the data (**Figure S2D**). Using a cell type decomposition analysis with CIBERSORTx^38^ (see Methods), we detected that this period of development coincides with the emergence of neurons (mean proportion of neurons ranging from 10% to 73% in the first (5-7PCW) and the last (14-17PCW) developmental group; **Figure S3A)**. Given these differences, we chose to account for this developmental trajectory in the joint analysis of the prenatal samples. The classification of the samples into discrete post-conception week categories, however, does not necessarily fully reflect the reality of development which is a continuous and complex process. Echoing this, we observed that using the developmental stages as a categorical variable in the DE analysis limits the power to detect sex-DE genes (only 31 sex-DE genes with q-value<0.01; **Figure S4A-B**) likely due to the low number of samples (2-39) per PCW for each sex.

Given the successful use of pseudotime inference in single-cell studies to model time course experiments^39^, we estimated a continuous pseudotime variable from the gene expression for each of the 266 prenatal RNA-seq samples using the phenopath R package^40^ (see Methods; **Figure S5A**). The inferred pseudotime variable correlated well with the developmental stages specified in the metadata (Kendall’s tau=0.46; **Figure 1B**), suggesting it accurately captures the trajectory of early brain development. We replicated this trend with pseudotime inferred by monocle2 (**Figure S5E-F**) and additionally confirmed that the inferred pseudotime was more consistent between samples from the same individual than between random samples (mean pseudotime standard deviation (SD) between samples from the same individuals = 0.30 vs mean pseudotime SD between random samples = 0.42, 1000 permutations, p-value=0.001; **Figure S5B**). Furthermore, we identified a clear correlation between cell type composition (obtained with CIBERSORTx; see Methods) and pseudotime (**Figure S5D**) in line with pseudotime tracking the developmental progression. For instance, the Neuronal Stem Cells (NSC) proportion decreased along pseudotime (Pearson’s r=-0.83, p-value=1.61x10^-69^) while the proportion of neurons (excitatory and inhibitory) increased (Pearson’s r=0.89, p-value=1.37x10^-91^). Overall, these findings show the inferred pseudotime variable reflects meaningful biology and allows ordering the samples along the development in a continuous manner.

### Widespread sex biases in the prenatal brain

We then carried out a sex-DE analysis across the 266 prenatal RNA-seq samples using a linear mixed model implemented in the variancePartition R package^41,42^ (see Methods). We found 3187 sex-DE genes (18.1% of all genes; q-value<0.01) (**Table S2**), with a slightly higher proportion of male-biased genes (N=1864, 58.5% of all sex-DE genes, binomial test p-value=8.96x10^-22^) (**Figure 1C**), observation in line with previous analyses of the developing brain^17,25^. Beyond the expected high number of sex-DE genes on the sex chromosomes (N=174, 28% of X and Y genes, after excluding the genes used for sample sex inference, see Methods), most of the sex-DE genes (94.5%) were autosomal. Only a few of the sex-DE genes (6 autosomal and 7 sex-chromosomal) showed substantial (|logFC|>1) effect sizes (**Figure 1C; Table S2**) with the average magnitude of differential expression between male and female samples being ∼20% (mean |logFC|=0.28). Of note, the sex-DE genes were on average more highly expressed and less constrained than the non-sex-DE ones (**Figure S6**).

For validation, we assessed sex-DE replacing the pseudotime variable with the reported developmental stages treated as a categorical variable. Reassuringly, all sex-DE genes (N=31) found in this analysis were also significant in the original analysis, and the high correlation of the effect sizes (logFC) between the two sex-DE analyses (Pearson’s r = 0.838 across all significant genes in the analysis with the pseudotime covariate, p-value<1x10^-10^) further indicated these two approaches are capturing the same sex-DE signal (**Figure S4B-C**) albeit with the analysis with the pseudotime covariate providing a more powered approach.

A Gene Ontology (GO) term enrichment analysis (see Methods) found that the female-biased genes were enriched in terms associated with the cell cycle (144/184 significant GO terms; **Table S3**), potentially related to neurogenesis^43^, while male-biased genes showed enrichment in biological processes related to mitochondrial metabolism and autophagy (39/115 significant GO terms; **Table S3**) that have been shown to be important in neuronal development^44,45^. Further pointing to the involvement of the genes in brain-relevant processes, male-biased genes were also significantly enriched in GO terms linked to synapses and neurons (3/115 significant GO terms).

### Systematic sex differences in the developmental trajectory of the brain

To complement the prenatal sex-DE analyses, we assessed differential expression for the pseudotime variable, i.e., a proxy for the developmental stage. Expectedly, illustrating the dynamics of gene expression during early brain development, a large number of genes (13129, 84.1%, after excluding the 3000 genes used to infer the pseudotime) were significantly DE (q-value<0.01) along pseudotime (referred to as pseudotime-DE genes) (**Figure 1D; Table S2**).

A joint examination of the two DE results revealed an enrichment in the overlap between the sex-DE and pseudotime-DE genes (2659/2756 genes, 96.5%; hypergeometric test, p-value=1.4x10^-39^ for enrichment) (**Figure 1E**), with considerable systematic consistency in effect directions in the two DE analyses. Most male-biased genes were upregulated along development, i.e., more highly expressed in later pseudotime (1592/1649 genes, 96.5%, p-value<1x10^-100^), while most female-biased genes were downregulated with pseudotime (1047/1107 genes, 94.6%, p-value<1x10^-100^) (**Figure 1E**). This observation was not explained by sex differences in the mean or range in pseudotime (mean male=0.06 [σ^2^=0.25], mean female=-0.02 [σ^2^=0.18]; Wilcoxon test p-value = 0.26; F-test p-value = 0.09; **Figure S5C**). Neither it was explained by a bias introduced at the pseudotime inference step (**Document S1**). Following the large overlap of these two gene sets, similar GO terms enriched in sex-DE genes were significantly enriched among the pseudotime-DE genes (**Table S3**). Interestingly, we detected only a few genes (496, 3.2%; **Table S2**) that had a significant interaction (q-value<0.01) between sex and pseudotime. This suggests that for most genes the magnitude of gene expression difference between males and females remains on average constant along pseudotime (e.g., *ZFX* in **Figure 1F**) irrespective of the large cell type compositional changes occurring in this developmental window (**Figure S3C**). This is in agreement with a previous study on developing brain reporting consistent age effects on gene expression in both sexes^27^.

### Attenuated sex bias in the adult brain

To understand the lifespan dynamics of the transcriptomic sex biases in the human brain, we additionally assessed the differential gene expression in 1633 adult (20 to 79 years; **Figure S7A**) forebrain samples from the GTEx project^35^. Following the analysis of the prenatal forebrain, we estimated the pseudotime for each adult sample before carrying out the differential expression analysis. Of note, the pseudotime in adult forebrain did not correlate with the age of the samples to a similarly high degree as in the prenatal forebrain (Kendall’s tau=0.12, **Figure S7B**), nor did pseudotime explain as large a fraction of the variability in the adult forebrain data (mean variance explained by pseudotime in the prenatal forebrain=16.4%; in the adult forebrain=3.4%; **Figure S8**). These observations likely point to a larger environmental variability in the adult samples compared to the prenatal samples. We detected 1033 sex-DE genes (6.0% of all genes; 91.4% autosomal) in the adult forebrain (q-value<0.01) (**Figure S7C**; **Table S4**), a considerably smaller number to prenatal forebrain despite the larger sample size of GTEx. Reflecting the lower number of sex-DE genes, the effect sizes of these genes in the adult brain were globally smaller than in the prenatal brain (sex-biased genes mean |logFC| 0.14 vs 0.28). Similarly to the prenatal brain, the male-biased genes were enriched in brain-related GO terms (e.g. regulation of neurotransmitters and synaptic organization) (**Table S3**).

When assessing DE along pseudotime in the adult brain, we identified 4549 (29.7%) differentially expressed genes (**Figure S7D**; **Table S4**). These genes were implicated in diverse processes such as synapse organization and autophagy for downregulated and upregulated genes, respectively (**Table S3**). The overall overlap of the sex-DE genes with the pseudotime-DE genes was significant (434/926 genes, 46.9%; hypergeometric test, p-value=1.63x10^-29^ for enrichment; **Figure S7E**), and again the sharing of effect directions was non-random. As opposed to the prenatal brain, in GTEx the male-biased genes were preferentially downregulated along pseudotime (283/610, p-value=2.0x10^-104^), and the female-biased genes enriched in the upregulated ones (104/316, p-value=3.1x10^-11^) (**Figure S7E**). Similarly to the prenatal brain, only a small number of genes (278, 1.8%) had a significant interaction (q-value<0.01) between sex and pseudotime (**Table S4**).

### Considerable sharing of brain sex bias between early development and adulthood

Gene regulatory mechanisms is known to change across development and life stages^46^. To understand what information is lost when using the more widely available adult tissue as a proxy for sex differences that occur during development, we applied several complementary strategies to estimate the sharing of sex-DE between adult and prenatal brain (**Figure 2**, **Table S5**). We show that the sex-DE between these two life stages are widely similar (72.2% consistency of effect size, Pearson’s r=0.60, π_1_=0.46; **Table S5**; **Figure 2A-B**), but still a degree lower than when comparing two independent datasets of the same tissue and age range (95.7% consistency of effect size, Pearson’s r=0.99, π_1_=0.87; **Table S5; Figure S9**) (see **Document S1**), suggesting also a degree of distinct sex effects between the two life stages.

**Figure 2:**
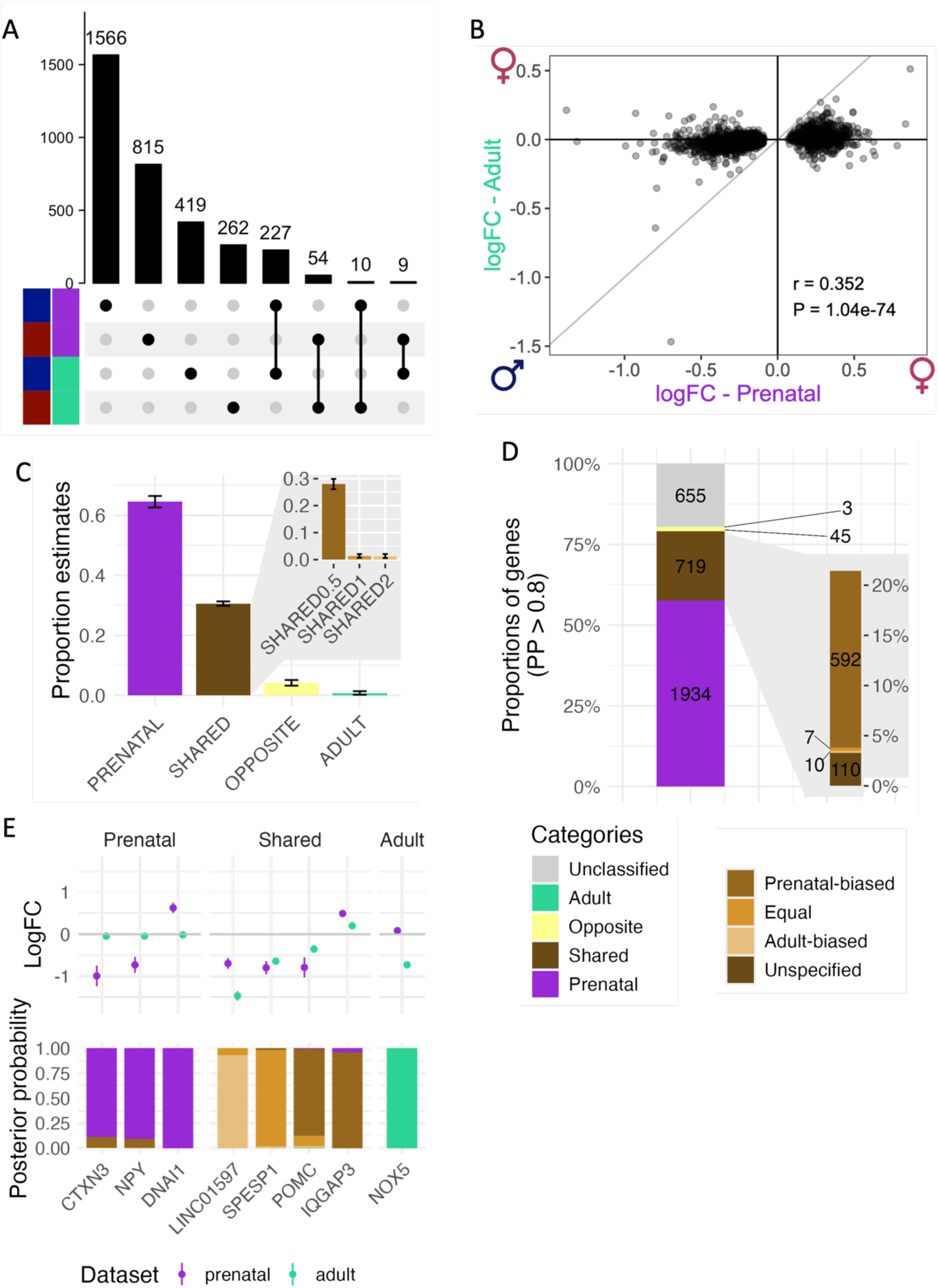
Comparison of the detected sex-DE genes in the prenatal (HDBR) and the adult (GTEx) forebrain. (A) Intersection of the sex-DE genes in the two datasets separated by the direction of effect (male or female-biased) as an upset plot. The upper barplot shows the size of the intersections. (B) Scatter plot of female/male effect size (logFC) in prenatal and adult dataset for all autosomal prenatal sex-DE genes. The Pearson’s correlation (r) of the effect sized and p-value associated (P) are given in the plot. (C) Proportion estimates and SE of each gene category in the Bayesian analysis: prenatal-specific (PRENATAL), shared (SHARED), opposite effect size (OPPOSITE) and adult-specific (ADULT). The zoom plot on the top right shows the proportion and SE of the different categories of shared effect: shared with an effect size twice larger in prenatal data (SHARED0.5), shared with the same effect size (SHARED1) and shared with an effect size twice larger in adult brain (SHARED2). (D) Proportions and numbers of individual sex-DE genes by effect category (Prenatal-specific, Adult-specific, Shared, Opposite and Unclassified). A gene was assigned into a category if the posterior probability for a given category reached >0.8. A zoom of the different shared categories (prenatal-biased, equal, adult-biased and unspecified) is shown on the right. (E) Examples of autosomal genes for the prenatal-specific (N=4), the shared (N=4) and the adult-specific (N=1) categories. The upper plot represents the logFC (with standard error) in both datasets. The lower plot shows the posterior probabilities from the Bayesian model for each gene.

To further quantify the life-stage specificity of sex-DE, we applied a Bayesian model comparison framework^47^ to estimate whether the sex-DE effect sizes are consistent with shared, opposite, prenatal-specific, or adult-specific patterns (see Methods; **Figure S10**). We analyzed the union of the 3356 sex-DE genes discovered in the two analyses (297 shared genes, i.e., significant sex-DE in both datasets, 2380 and 679 significant only in the prenatal and adult data, respectively) using this approach.

In this analysis, two major groups of sex-DE appeared, covering more than 95% of the detected DE effects. Most of the sex-DE signal, 64.6%, was assigned to prenatal-specificity, in line with most of the sex-DE genes originating from the prenatal analysis (i.e. 70.9% of the 3356 input genes were only sex-DE in the prenatal dataset) (**Figure 2C**). However, the shared male/female differences between the two datasets accounted for 30.6% of the effects in total, a considerably greater proportion than estimated based on significant DE genes only (8.3% in the input genes with the same direction of effect). This is, nevertheless, a degree lower than the 64.3% of sex-DE sharing for the same tissue of different individuals (adult cortex, GTEx vs BrainSeq) (**Figure S9C**). The great majority of the shared signal (28.1% of all sex-DE genes) was assigned to the model supporting shared effects with a magnitude of effect twice larger in the prenatal brain than in the adult brain (SHARED0.5) (**Figure 2C**) aligned with the earlier observation of larger sex-DE effect sizes in the prenatal brain.

Only small fractions of the genes were found to have opposite sex-DE effects (of same magnitude) between the two life stages (4.2%) and adult-specific sex-DE (0.7%). The latter finding is striking considering that, 20.2% of the input genes were significantly DE only in the adult brain. However, many of these adult sex-DE genes showed enrichment for small p-values in prenatal data (π_1_=0.56) but failed to reach significance (see Document S1), which explains why the model classified them as shared genes. Given these observations, it appears there is very little sex-biased expression captured exclusively in the adult brain but that the early developing brain already displays a considerable fraction of sex-DE that is maintained into adulthood albeit with a typically lower degree.

### Shared sex differences in prenatal and adult brain reflect neuronal functions

To allow for the examination of characteristics defining the shared and developmental stage-specific sex-DE, we compiled sets of individual genes that were confidently assigned to one of the sex-DE patterns in the model comparison framework. To this end, we included all genes that had a gene-specific posterior probability (PP) larger than 0.8 for one of the four possible categories of sex-DE effects across the prenatal and adult data sets (see Methods; **Figure 2D**). This approach resulted in the classification of 1934 genes into the category of prenatal-specific effect (e.g. *DNAI1*, *NPY*, *CTXN3* genes; **Figure 2E**), 719 into shared effect (a sum of PPs of the three categories of shared effects) (e.g. *IQGAP3*, *POMC*, *SPESP1*, *LINC01597*; **Figure 2E**), the majority (82.3%, 592/719) of which had a larger effect size in the prenatal forebrain (PP.SHARED0.5>0.8), 3 into adult-specific effect (e.g. *NOX5*; **Figure 2E**), and 45 into the opposite effect (**Table S6**; **Figure 2D**). 655 sex-DE genes were left unclassified as they did not reach the chosen threshold (PP>0.8) for any of the models, partly driven by their lower sex-DE effects in prenatal brain (**Figure S11**). As the shared and prenatal-specific sex-DE genes constituted the two largest categories, in the following sections, we focused on characterizing these two sets further. Of note, as before, the full set of 3356 sex-DE genes tended to be less constrained compared to non-sex-DE genes (Wilcoxon test p-value=5.3x10^-08^; **Figure S12A**).

We first set out to explain the high degree of shared DE between the prenatal and adult brain, and therefore the lack of adult-specific DE, by investigating their gene expression characteristics and gene functions. While the sex-DE genes were generally more highly expressed in the brain than genes without sex differences (Wilcoxon test p-value=2.2x10^-03^ and 1.9x10^-19^ in prenatal and adult data respectively; **Figure S12C-D**), the two subsets displayed differences in their patterns of gene expression. The prenatal-specific sex-DE genes were expectedly more highly expressed in the prenatal than in the adult brain (Wilcoxon test p-value=8.1x10^-11^) (**Figure S13A**). The opposite was observed for the shared sex-DE genes that displayed higher expression in the adult brain (Wilcoxon test p-value=4.0x10^-04^) (**Figure S13A**). This suggests that the consistency of the level of expression in the bulk tissue sample does not explain the shared sex-DE pattern. However, as the cell type proportions between the developing and adult brain differ considerably (**Figure S3C-D**), the bulk tissue-level expression level may not show the patterns associated with individual brain cell types.

We therefore investigated whether the shared sex-DE signal might be driven by specific cell types shared but present at different proportions between the prenatal and adult brain. We tested this by comparing the level of expression of these two gene lists in different cell types from the human brain (single-cell RNA-seq data from ^48^; see Methods). The shared sex-DE genes were more highly expressed in neurons (Wilcoxon test p-value adjusted for multiple testing=6.4x10^-04^ and 1.9x10^-03^ for excitatory and inhibitory neurons respectively) (**Figure 3A**) than the prenatal-specific sex-DE genes. The neuron-enriched expression provides a plausible explanation of their consistency across the two studied life stages. In contrast, the prenatal-specific sex-DE genes appeared more shared across different cell-types, as they were more highly expressed in progenitor cells (Wilcoxon test p-value adjusted=4.9x10^-07^ and 1.4x10^-03^ in neuronal and oligodendrocyte progenitor cells respectively) (**Figure 3A**) and other non-neuronal cells (Wilcoxon test p-value adjusted=3.9x10^-08^, 4.3x10^-07^, 8.3x10^-07^, 8.3x10^-07^ and 4.7x10^-07^ in oligodendrocytes, astrocytes, pericytes, microglia and endothelial cells respectively) (**Figure S13B)** compared to the shared sex-DE genes.

**Figure 3:**
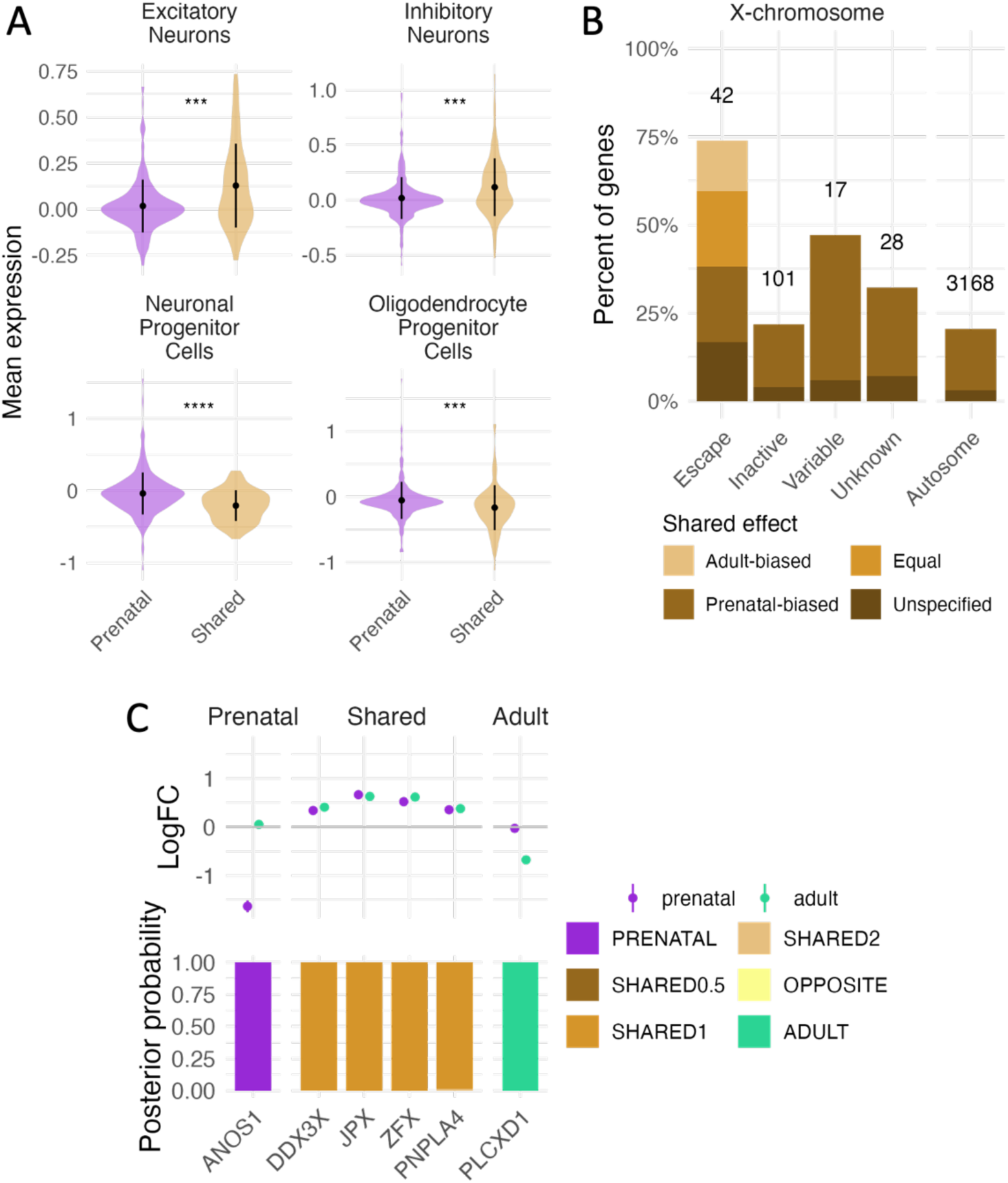
Characteristics of the shared and prenatal-specific sex-DE genes. (A) Expression of the shared and prenatal-specific genes in excitatory neurons, inhibitory neurons, neuronal progenitor cells and oligodendrocyte cells. Difference in mean expression between the two categories was tested with a Wilcoxon test. *** and **** indicate Wilcoxon test p-value p <= 0.001 and <= 0.0001 respectively. (B) Percentages of genes in each shared category (shared with the same or different effect size) for different classes of X chromosome inactivation (XCI) genes (escape, inactive, variable or unknown) and autosomal genes. The number of genes per category is shown as the top of each bar. (C) Examples of XCI escape genes classified as prenatal-specific (N=1), shared (N=4) or adult-specific (N=1). The upper plot represents the logFC (with standard error) in both datasets. The lower plot shows the posterior probabilities from the Bayesian model for each gene.

In agreement with the above findings from cell type-specific expression, the prenatal-specific sex-DE genes were also more broadly expressed across adult human tissues from GTEx than the shared sex-DE genes (Wilcoxon test p-value=6.7x10^-27^) (**Figure S13C**) suggesting a larger diversity of functions of these genes. Further, aligned with the neuron-specificity of the shared sex-DE genes, the analysis of tissue specificity of expression across the GTEx tissues for these same set of genes confirmed that the shared sex-DE genes were more tissue specific than the prenatal-specific sex-DE ones (Wilcoxon test p-value=9.0x10^-36^) (**Figure S13D**).

The GO analyses further indicated neuronal functions contributing to the shared sex-DE between the prenatal and adult brain (**Table S3**). While the shared female-biased genes were not enriched in any GO term, the shared sex-DE genes with male-biased expression displayed enrichments in terms linked to neuron and synapse activity (45/146 significant GO terms). These enrichments were distinct from the ones detected for the prenatal-specific sex-DE genes but showed overlap with the enrichments discovered for the adult brain sex-DE. The prenatal-specific sex-DE genes recapitulated similar GO enrichments to the full prenatal sex-DE results (**Table S3**) with the addition that female-biased genes were enriched in GO terms related to development (17/172), potentially pointing to relevance for sex biases in brain development (**Table S3**).

### Early prenatal testosterone surge as potential driver of prenatal-specific expression sex differences

Given that most of the identified sex-DE genes were specific to the prenatal life stage, we additionally set out to understand the biological processes influenced by these sex differences in gene expression and the plausible underlying mechanisms. Since the studied time span coincides with the first testosterone surge (proposed to occur between 9 and 18 PCW^49,50^), we hypothesized that hormonal factors could partly contribute to the expression of these early development specific sex-DE genes. To test for this, we analyzed the transcription factor binding sites (TFBS) enriched in these genes’ promoters using the Unibind framework^51^ (see Methods).

While no TFBS were significantly enriched (q-value<0.05) in the genes that showed shared sex-DE throughout lifespan, the prenatal-specific genes displayed several enrichments, with female-biased genes enriched in 8 TFBS, and the male-biased ones in 74 TFBS. Notably, these enrichments included established hormonal transcription regulators like androgen receptor (*AR*) (q-value=2.7x10^-4^, enrichment odds ratio in male-biased genes=1.5) and Estrogen Receptor 1 (*ESR1*) (q-value=0.03, enrichment odds ratio in male-biased genes=1.3) (**Table S7**). Moreover, some of the 74 TFBS enriched for male-biased genes were known *AR* interactors, like *FOXA1* and *FOXP1*^52^, or *ESR1* interactors, like *JUN* (from the Reactome database^53^). Interestingly, *AR* expression was female-biased only in the prenatal brain (prenatal logFC=0.32; adult logFC=0.05). This up-regulation of *AR* in females compared to males is in line with the known auto-downregulation of *AR* in presence of androgens^54^.

The enrichment of prenatal-specific sex-DE genes in androgen-responsive genes was further validated using a set of genes that have been shown to be differentially regulated with androgen treatment^55^ (**Figure S14**). We observed a more than two-fold enrichment of female-biased genes in genes up-regulated with testosterone treatment in NSCs (relative enrichment=2.61, hypergeometric test p-value=4.20x10^-08^), while the prenatal-specific male-biased genes were enriched in genes down-regulated by the same testosterone treatment (relative enrichment=1.85, hypergeometric test p-value=9.75x10^-23^). Similar enrichment patterns were observed for other androgen treatments (e.g. DHT). In contrast, the shared sex-DE genes were not significantly enriched in genes regulated by these androgen treatments, pointing to the specificity of the hormonal effects to the prenatal stage. We propose that these hormonal effects may be related to the early testosterone surge.

### Lifelong gene expression sex bias due to escape from X chromosome inactivation

To assess other potential processes contributing to the sharing of sex-DE, we investigated the role of X chromosome inactivation (XCI), prompted by the higher proportion of X-chromosomal sex-DE genes (proportion test p-value=2.22x10^-06^) among the shared sex-DE genes. Given the known early establishment of XCI^56^ and the reported tissue similarities in the sex-biased expression in adulthood introduced by escape from XCI^57^, we reasoned that the X-chromosomal genes that escape from XCI would display high sharing of sex-biased expression also across life stages. Indeed, known XCI escapees (N=42 with sex-DE, 22.2% of X-chromosomal sex-DE genes) displayed considerable concordance in sex-DE in the prenatal and adult brain in comparison to autosomal (N=3168) and inactive X-linked genes (N=101). Altogether, 73.8% (31/42) of the escape genes were classified as shared sex-DE genes compared to 21.8% (22/101) of known inactive X-chromosomal genes (ꭓ^2^ p-value = 1.37x10^-08^ for difference in proportions) (**Figure 3B**). Further, unlike most other shared sex-DE genes, the XCI escapees were typically more consistent in their degree of sex bias between the two life stages. Altogether, 21.4% (9/42) of the escape genes (vs. 0/101 inactive genes, p-value=9.52x10^-06^) displayed similar effect sizes in the prenatal and adult brain (PP.Shared1>0.8), including *DDX3X* (PP.Shared1=0.997), *JPX* (PP.Shared1=1.000) and *PNPLA4* (PP.Shared1=0.986) with large female-biased effects (logFC range=0.34-0.66, corresponding to 27-57% higher expression in females) (**Figure 3C**). These observations suggest escape from XCI is a highly stably maintained phenomenon irrespective of changes in the environment or cell type composition.

Interestingly, *ZFX*, a known XCI escapee and a transcription factor proposed to influence autosomal sex-DE^58^, was consistently female-biased across the developing and adult brain (prenatal logFC=0.52, adult logFC=0.62, PP.Shared1=1). Supporting the role in autosomal gene regulation, *ZFX* binding sites were enriched in the prenatal male-biased genes (q-value=0.02, enrichment odds ratio=1.3; **Table S7**), but the enrichment did not reach significance when focusing on either the female-biased genes or shared and prenatal-specific sex-biased genes only. The enrichment of *ZFX*-regulated genes among the male-biased genes, rather than the female-biased ones, could be explained by *ZFY*, a homologous gene in the male-specific region of the Y chromosome (highly male-biased, prenatal logFC=-8.69 and adult logFC=-8.43), known to bind to similar motifs and occupy similar genomic locations than *ZFX*^58^.

Despite this general consistency across the life stages, a few escapees (N=11 with PP.Shared<0.8) displayed more divergent patterns of sex-DE. For instance, *PLCXD1*, a gene in the pseudo-autosomal region of the X chromosome, was the only escape gene with evidence for adult-specificity of sex-DE with large male-biased effects exclusively in the adult brain (logFC=-0.68, PP.Adult=1.00) (**Figure 3C**). Also, four escape genes displayed evidence for prenatal specificity. These include *ANOS1* that was highly male-biased in expression, an unusual pattern for an escape gene in the non-pseudo-autosomal region^57^, in the prenatal brain (logFC=-1.63) and non-significant in the adult brain (logFC=0.054, PP.Prenatal=1.00) (**Figure 3C**). *ANOS1* is crucial for neuronal migration in the developing brain^59^ and was previously suggested as a gene subject to variable escape in adult human tissues^57^. In general, these variable patterns of escape can point to cell-type dependent control of XCI but can also reflect, e.g., hormonal or other age-related regulation of gene expression dampening the expected sex bias arising from escape. Further, 25 previously established escape genes did not show significant sex bias in either of the life stages, possibly due to their lower level of expression in the brain (**Figure S15A**) but this finding could also suggest a lower level of escape not sufficient to result in significant sex biases in expression. Supporting this latter hypothesis, these genes were also less frequently observed as sex-biased in diverse adult human tissues than other escape genes (**Figure S15B**).

### Genes with shared sex bias associate with disease networks

As the final step, we set out to understand the potential phenotypic relevance of these lifelong and life stage-specific sex biases in the forebrain gene expression. To this end, we examined the enrichment in overlap of the observed sex-DE patterns with genes implicated in the etiology of brain disorders identified through both exome sequencing and genome wide association studies (GWAS).

While we found earlier that the sex-DE genes display roles in central brain-related processes, across the 19 unique diseases studied we, however, found little conclusive evidence for enrichment in overlap of sex-DE and brain disease-associated genes in any of the sex-DE gene lists (**Figure S16A-B**; **Figure S17A-B**; **Table S8**). For the 9 gene lists derived from exome sequencing studies, these generally representing high-impact disease genes where the functional loss of another gene copy increases in the disease risk, we observed no significant enrichment (hypergeometric test p-value < 6.9x10^-04^ = 0.05/9x8) with the prenatal-specific or shared sex-DE genes. Four disease gene lists displayed a nominally significant enrichment for sex-DE and these findings were further validated by a comparison against a background of genes with a similarly high expression level in the brain (see Methods). For instance, genes implicated in neurodevelopmental disorders (NDD, 662 genes) appeared enriched among the female-biased shared sex-DE genes (1.6 relative enrichment, hypergeometric p-value = 0.003, permuted p-value for enrichment < 0.001) (**Figure S17A**), and genes implicated in schizophrenia (244 genes) showed an enrichment among the male-biased shared sex-DE genes (2.3 relative enrichment, hypergeometric p-value = 0.029, permuted p-value = 0.004). The overlap with GWAS associations, where the association is typically mediated through regulatory effects, similarly found no significant enrichments (MAGMA p-value < 6.9x10^-04^ = 0.05/9x8) in small GWAS p-values at and near the sex-DE genes. Some of the nominally significant findings, nevertheless, corroborated the earlier results, e.g., male-biased shared sex-DE genes were enriched in schizophrenia associations (MAGMA p-value = 0.012). Such limited direct disease overlap can be at least partly expected, as genes with phenotypic effects tend to tolerate less genetic and expression variation^60–62^. In line with these observations, we noted that the identified sex-DE genes, in general, were less constrained than genes without sex differences (**Figure S6B; Figure S12A**). A similar lack of significant enrichment was observed when focusing on the most constrained genes (i.e. 3 lowest constraint bins; **Figure S12B**).

In the light of these findings, we hypothesized that the sex-DE genes could exert phenotypic effects through being interaction partners to disease-associated genes, rather than the disease genes themselves being directly impacted by sex-biased expression. To test for the presence of such more distal effects, we used the brain-specific gene co-regulation networks from the MetaBrain resource^63^, constructed of >8000 harmonized adult brain RNA-seq samples. This network has been applied in prioritizing genes linked with genes in associated GWAS loci for brain-related conditions, which has pinpointed, e.g., 2737 genes (5% false discovery rate) highly likely to be co-regulated with genes within the schizophrenia associated loci. Here, we observed that the shared female-biased sex-DE genes were significantly enriched (hypergeometric test p-value < 7.8x10^-04^=0.05/8x4 and permuted p-value<0.05) in the cortex-specific co-regulatory networks of both schizophrenia (**Figure 4**) and multiple sclerosis (MS) GWAS genes (**Figure S17C**; **Table S8**) (p-value = 3.21x10^-17^ and 1.36x10^-06^, respectively). Additionally, the adult sex-DE genes were significantly enriched in the co-regulatory network of MS GWAS genes (**Figure S16C**). The association with schizophrenia was not driven by the GWAS genes themselves, as no enrichment was detected in overlap with schizophrenia GWAS genes from MetaBrain (e.g. for shared male-biased genes, p-value = 0.06). While the direction of the sex bias effect for a co-regulation network is difficult to translate to the direction of the impact on the phenotype, these findings, nevertheless, suggest that a possible phenotypic relevance of the brain gene expression sex biases may arise through the modulation of the activity of the gene networks involved in mediating the genetic disease risk.

**Figure 4:**
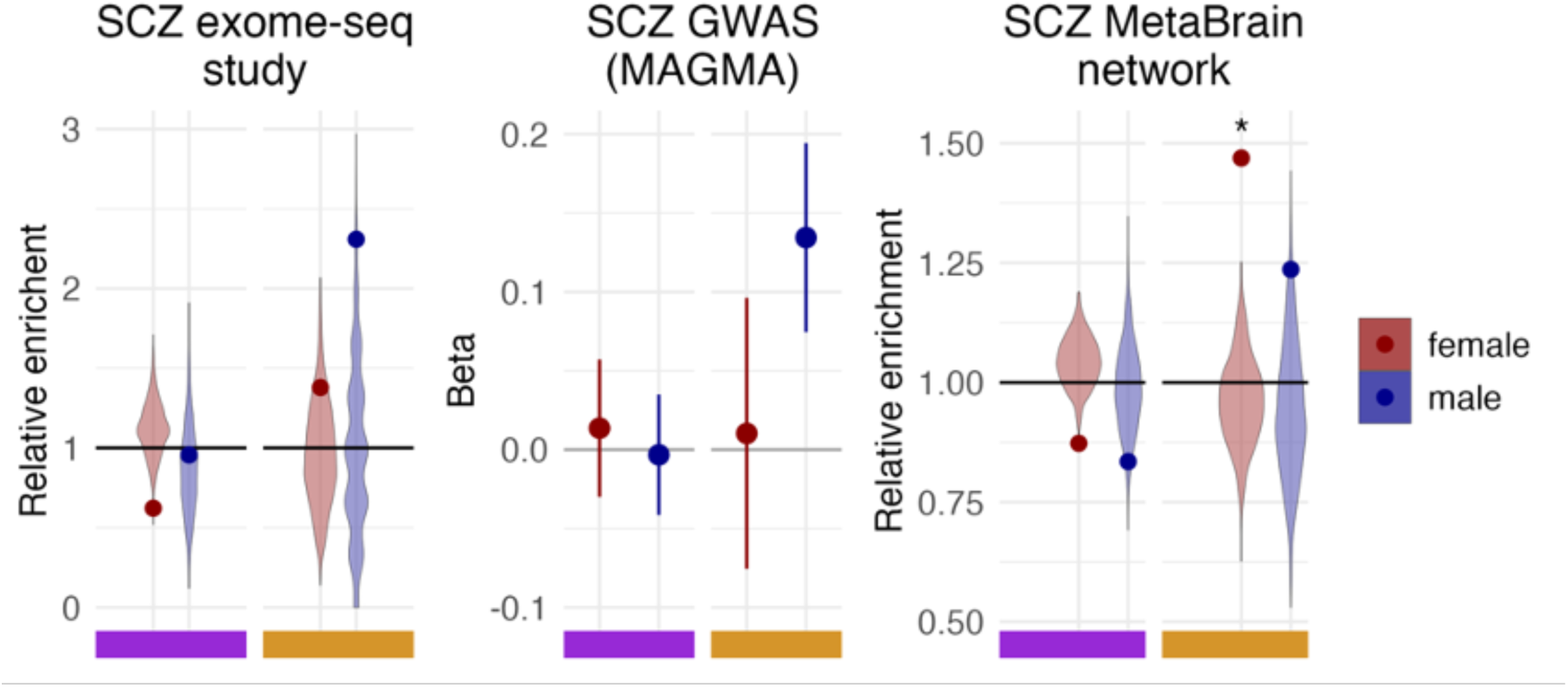
Enrichment analysis of three different schizophrenia gene lists in the shared and prenatal-specific sex-DE genes. (left) Genes from an exome-sequencing study of SCZ (N=244). (middle) Genes from a SCZ GWAS analyzed with MAGMA. (right) Genes from a SCZ co-regulation network analysis from the MetaBrain resource (N=2737). The colored box at the bottom indicates from which data the results are (violet = prenatal and yellow = shared). The violinplots on the left and right plots represent the distribution of the relative enrichment for the 1000 random gene lists and the point the enrichment for the true gene list. The error bars in the middle plot depict the SE of the beta values. * denotes significant enrichment of the SCZ gene list

## Discussion

To better understand the mechanisms underlying the widespread sex biases observed in brain structure, function, and various brain-related conditions, we set out to characterize sex differences in the human brain transcriptome during early development and adulthood. To this end, we used data from the HDBR, that spans the period of neurogenesis in prenatal development (5-17 PCWs), and from adult brain samples from the GTEx project, and took advantage of statistical methods to account for the high transcriptional variability and to allow for robust comparisons of patterns of sex-DE between the prenatal and adult brain. We discovered that transcriptional sex differences are widespread and systematically present already in the early brain development and further show that a large fraction of these sex biases are maintained into adulthood in a manner where most of the adult brain sex differences are already present in the prenatal brain. Overall, our work demonstrates the relevance of the fetal development to sex differences and highlights the roles of sex-chromosomal and early hormonal influences in shaping sex-biased brain biology.

While sex biases in the adult brain have been extensively studied, both in terms of gene expression and its genetic regulation^16,64^, the scarcity of early brain samples and the inherent heterogeneity of the available datasets has posed limitations to the study of sex differences the fetal brain. In our analyses, we took advantage of the brain expression data from the unique HDBR resource^34^, a collection of early embryonic and fetal brain samples, applied also in previous investigations into prenatal sex differences^25,26^. To facilitate the discovery of sex-DE in these prenatal brain samples, we accounted for the high transcriptional variability by focusing our assessments on the forebrain region and by additionally modelling the active developmental period using pseudotime, which allowed us to better align the samples across the developmental time points than the ordinal categories typically used. With this approach, our analyses of the HDBR samples revealed an abundance of sex-DE genes during the very early development. Up to 18% of the developing forebrain transcriptome displaying significant (q-value<0.01) male-female differences, that reflect relevant brain development-related processes, including cell cycle^43^, mitochondrial metabolism^44^ and autophagy^65^. Together, these findings provide support for the early origins of brain sex differences.

Although abundant, the detected sex differences in gene expression were mostly small in scale, aligned with findings from adult tissue^16^. The average sex difference in gene expression of around ∼20% is comparable to the allelic effects of common gene regulatory variants^66^, and only a few genes showed sex differences similar to those observed in pathological conditions (e.g. 1567 DE genes with |logFC|>1 in the cortex of ASD patients compared to healthy individuals^67^). A likely implication of this finding is that the developing brain tolerates only a limited amount of variability between groups of normally developing individuals. This is further supported by our finding that sex-DE typically impacts genes that are less constrained, i.e., genes that are more tolerant to genetic variation, suggesting that many crucial brain processes are in fact shielded from expression variation between the sexes.

To understand the dynamics of brain sex biases across life, we compared the prenatal sex-DE to that discovered in the adult forebrain samples from GTEx. Through the application of a Bayesian model comparison framework on the identified sex-DE genes, we found two dominant patterns of sex-DE. Both the high fraction of prenatal-specific sex-DE (64%) and the substantial sharing of sex-DE between prenatal and adult forebrain (30%), despite the considerable differences in gene expression and cell type composition, further point to the relevance of early development for brain sex biases. The apparent lack of adult-specific sex-DE in our analyses suggests the sex biases detected in the adult brain are actually largely emerging earlier in brain development. This finding of the limited adult-specificity contrasts with recent evidence from Rodríguez-Montes and colleagues^68^, where sexual maturity was found as a crucial time point for the onset of sex-biased expression. We attribute these differences in findings mostly to the larger sample size in the current study with respect to the fetal time point, and our approach that considers the similarity of sex effects beyond those genes that display significant sex differences at a given life stage. Similar approach found sharing but not to such a high degree between human and primate brain^69^. Indeed, if applying strict p-value cutoffs, only about a quarter of the adult sex-biased genes would be considered sex-DE in the prenatal brain. Our findings provide one potential mechanistic explanation for the large sharing of sex-DE whereby the consistency of sex-DE may be driven by the shared neuronal cell types between the prenatal and adult brain.

In contrast to later life stages, the effect of the environment is likely small in the very early stages of brain development, and as such we hypothesize that the sex biases in the fetal human brain primarily originate from intrinsic biological factors, such as direct sex chromosome effects or early hormonal influences^70^. Indeed, we found support for both processes shaping the transcriptional sex differences, partly in a life-stage dependent manner. Following the finding that sex-DE genes specific to the fetal stages are enriched for transcription factor binding sites of hormone receptors, we propose some of the observed sex effects are reflections of the significant hormonal changes occurring during this developmental window owing to the fetal testosterone surge (estimated to occur between 9 and 18 PCW^49,50^). Notably, only the prenatal-specific sex-DE genes, and not the shared sex-DE genes, exhibited effects associated with hormonal regulation. This suggests that the transient influence of the early testosterone surge extends beyond sex determination and genitalia formation to also impact sex-biased gene expression in the fetal brain, potentially leading to stable sex differences in the organization and function of the brain.

Sex-chromosomal contributions to sex-DE were evident in the clear sex biases of genes escaping from XCI, in line with earlier reports^16^. We further found that XCI escapees are a unique category of early sex-DE genes where the sex effects are highly consistent in magnitude between the prenatal and adult brain. This confirms the early establishment of XCI and escape^71,72^, adds to the known similarities in escapee expression sex biases in adult tissues^57^, and points to escape from XCI being a highly stably maintained phenomenon throughout life irrespective of cell type compositional, hormonal and other changes. As XCI escapees have proposed roles in the regulation of autosomal gene expression^58,73^ the substantial and consistent sex-DE associated with escape raises the possibility of escapees introducing broader sex effects in the brain beyond the sex-chromosomal expression.

Understanding how the life stage specific and shared sex effects in gene expression relate to phenotypic sex differences is a key open question. Despite the similar processes implicated by sex differential expression and genetic studies of neurodevelopmental disorders, we detected limited overlap between sex-DE and disease genes. Similar observations of the lack of overlap between fetal cortex sex-DE were recently made with regards to ASD risk genes^27^. We focused our assessments on genes linked to diseases through genetic studies (GWAS or exome sequencing) allowing us to examine genes with confident links to the disease etiology rather than genes identified through case-control comparisons of gene expression (e.g. ASD-related alterations^69^) that could rather reflect consequences of the disease process. We attribute the lack of overlap sex-DE and disease genes to systematic differences between sex-DE and GWAS signals, e.g., in terms of gene constraint, in analogy to recent findings on the distinct characteristics of common gene regulatory variation and GWAS hits^62^. In the light of our findings, we propose that the brain sex-DE may nevertheless play a role in differential disease susceptibility through impacting the networks the disease genes are involved in.

## Limitations of the study

In our study, we focused on comparing patterns of sex-biased gene expression from two life stages, the prenatal development and adulthood. Other time points across the development, e.g., childhood, adolescence, and puberty, would allow us to build a clearer picture of the life course dynamics of sex-DE. Further resolution for the very early developmental time points, before the first testosterone surge (before 9 PCW), would additionally be valuable for a more detailed evaluation of the onset of sex-DE and the dissection of the hormonal influences and the direct sex chromosome effects on early sex-DE. The sex-DE effect sizes in the adult brain tend to be smaller than in early development, which is a potential reflection of the increased expression variability associated with aging^74^, induced by the environmental influences that we are unable to fully account for in the analysis. This may complicate the inference of the proportions of genes that show sex-DE across lifespan, or only in adulthood, and lead to an underestimation of these proportions. Also, we currently lack an understanding of the potential consequences of the prenatal-specific sex-DE, and hence much more work is needed in this space. For instance, the study of the co-regulatory network genes might bias our findings toward adult and shared genes as the networks are constructed from adult tissue.

## Conclusions

We present a comprehensive evaluation of the emergence and dynamics of transcriptomic sex differences in the developing and adult forebrain. Our findings highlight the fetal development as a crucial time point for the introduction of brain-related sex differences and propose the involvement of both hormonal and sex-chromosomal contributions in shaping the brain sex biases. Evaluation of further time points in development and cell-type specific data may help in delineating the causes and consequences the transcriptomic sex biases in more detail. Although it remains unclear how the identified transcriptional sex biases are mechanistically linked with brain structure or function, or with neurodevelopmental phenotypes, our findings point to the direction that a degree of the sex differences in brain-related traits stem from non-environmental effects originating from the prenatal period. This can have implications for the understanding of sex-biased disease susceptibility and the development of more targeted diagnostics and therapeutics.

## Acknowledgments

T.T. was funded by the Sigrid Jusélius Foundation (https://sigridjuselius.fi/en/), the HiLIFE Fellows Program, and the Research Council of Finland (https://www.aka.fi/en/) grant numbers 315589 and 320129.

## Author contributions

The project was conceived and planned by C.B.P. and T.T. C.B.P. performed the analyses under the supervision of T.T. J.A. conducted the MAGMA analysis and J.K. preprocessed the prenatal dataset. C.B.P. and T.T. wrote the manuscript, with J.L., J.A., and M.J.D. providing comments and edits.

## Declaration of interests

M.J.D. is a founder of Maze Therapeutics.

All other authors declare no competing interests.

## Data availability

All raw data used in this article were previously generated by HDBR and GTEx, and are publicly available in the EBI ArrayExpress database under accession number E-MTAB-4840 and in the GTEx online portal https://www.gtexportal.org/home/ respectively. The processed data supporting the conclusions of this article are included in the supplementary tables. The codes used in this study are openly available at https://github.com/cbenoitp/sexDE_prenatal_brain.

## Supplemental information

### Supplemental document

**Document S1:** Supplementary results

### Supplemental tables

**Table S1:** Description of the datasets. Number of the samples used in this study by dataset, tissue, sex, and age, and scRNA-seq datasets used in the cell type decomposition analysis.

**Table S2:** Differential expression results in prenatal forebrain.

**Table S3:** GO term enrichment analysis results.

**Table S4:** Differential expression results in adult forebrain.

**Table S5:** Comparison metrics between sex-DE genes

**Table S6:** Bayesian model results for shared sex-DE effect between prenatal and adult forebrain.

**Table S7:** Transcription factor binding sites (TFBS) analysis results.

**Table S8:** Enrichment analysis results.

### Supplemental figures

**Figure S1:**
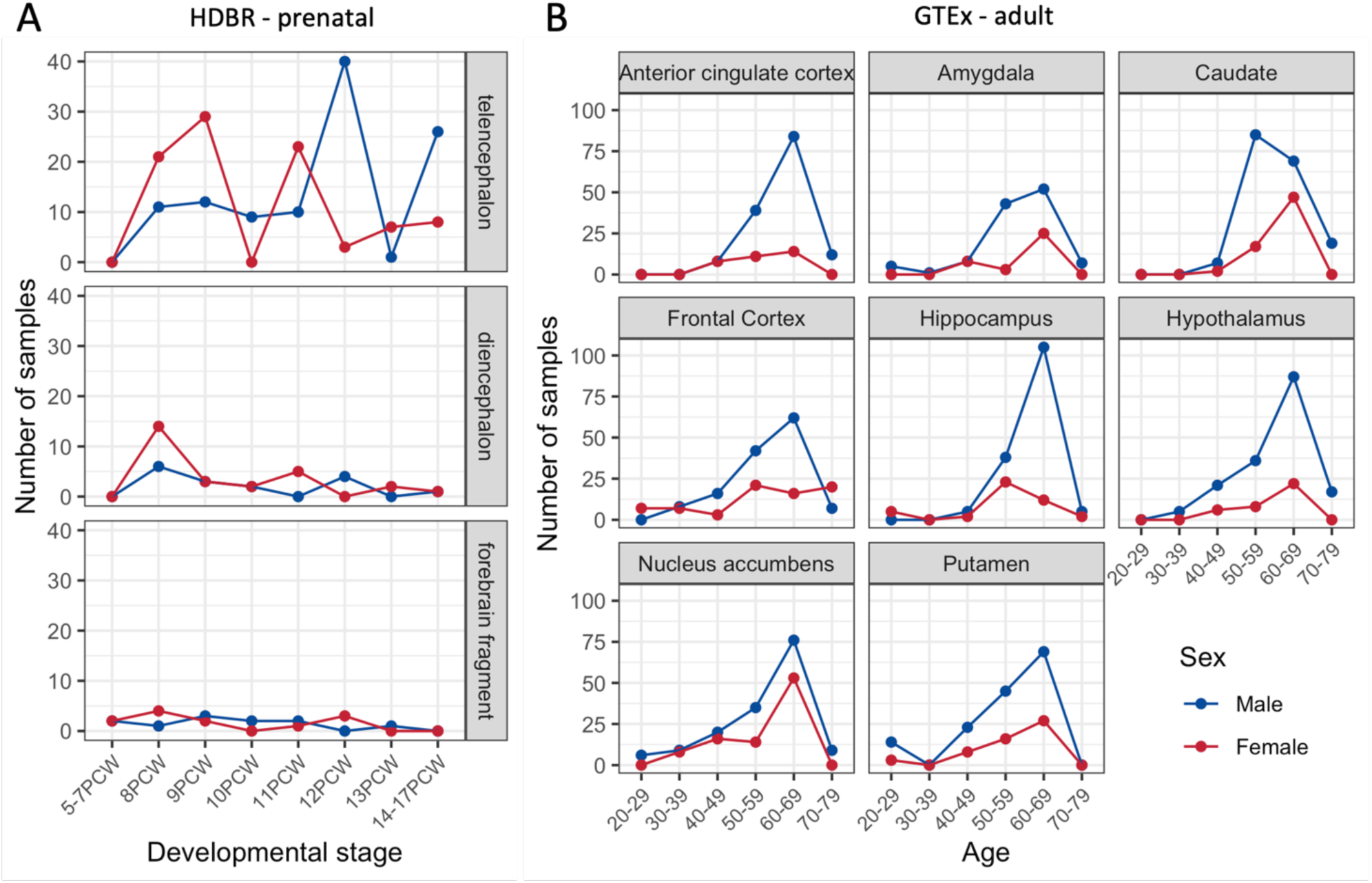
Number of samples per sex, developmental stages/ages and forebrain region in (A) HDBR and (B) GTEx.

**Figure S2:**
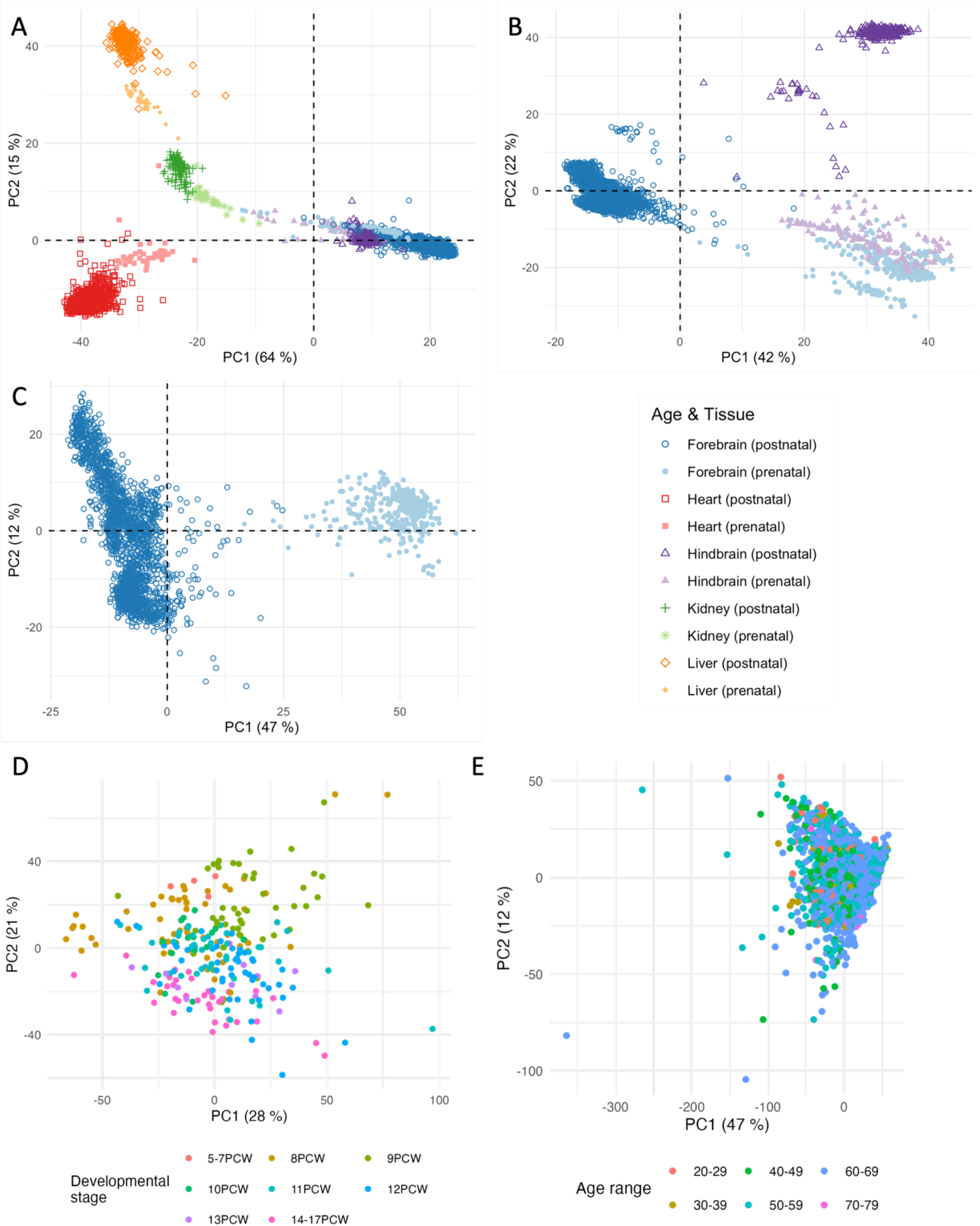
Principal component analyses (PCA) plots for prenatal and postnatal samples from (A) all tissues, (B) brain tissues and (C) forebrain only. Each sample is colored by tissue type and their shape correspond to the age (prenatal and postnatal). (D) PCA for prenatal forebrain samples from HDBR colored and shaped by developmental stage. (E) PCA for adult forebrain samples from GTEx colored by age.

**Figure S3:**
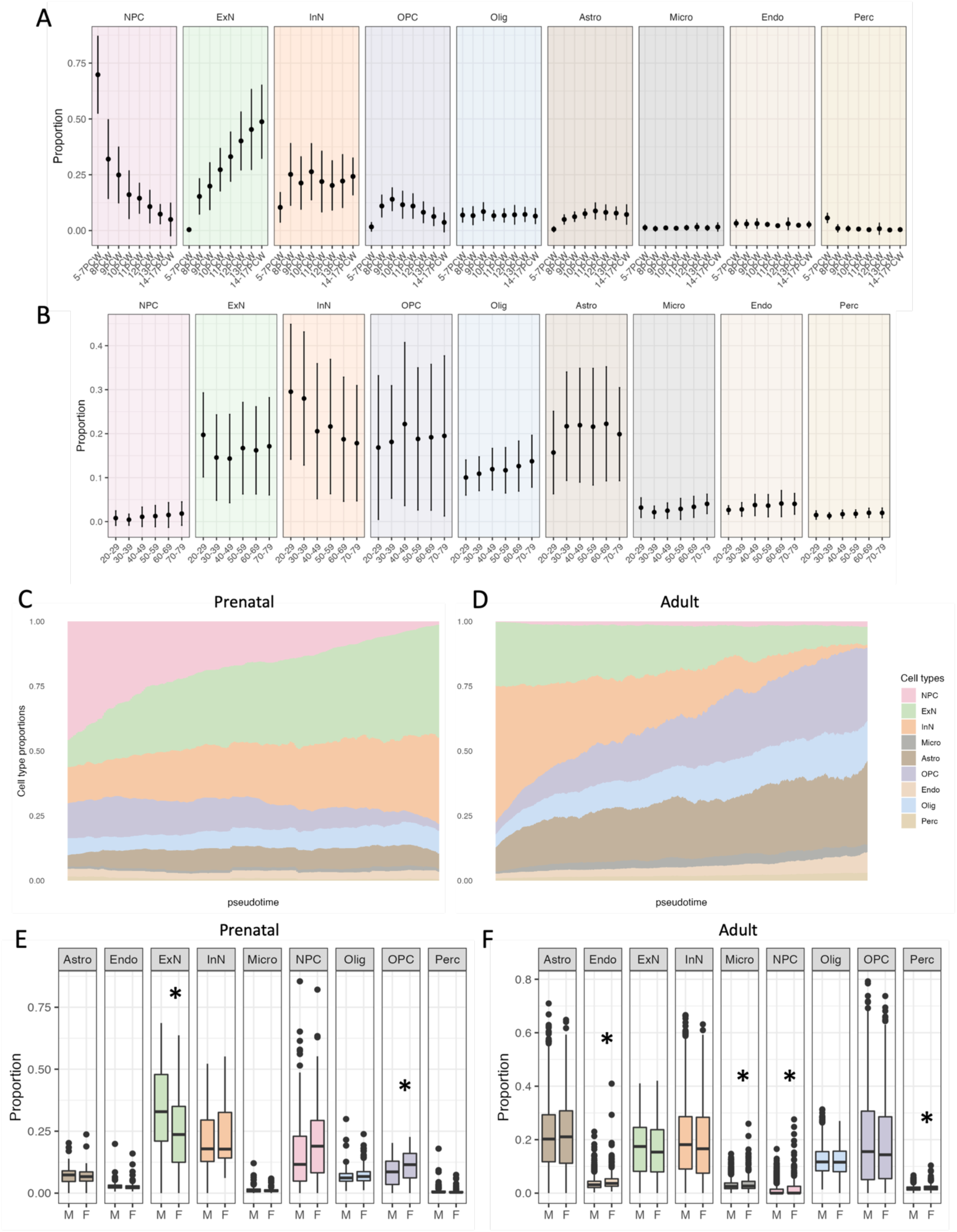
Cell type decomposition. (A) Proportion and SE of each cell type for the prenatal samples grouped by developmental stages. (B) Proportion and SE of each cell type for the adult samples grouped by age group. (C & D) Moving average along pseudotime of cell type composition for (C) prenatal and (D) adult samples. (E & F) Cell type proportion by sex for (E) prenatal and (F) adult samples. *: p-value adjusted < 0.05. NPC = Neural Progenitor Cells; ExN = Excitatory Neurons; InN = Inhibitory Neurons; OPC = Oligodendrocyte Progenitor Cells; Olig = Oligodendrocytes; Astro = Astrocytes; Micro = Microglial cells; Endo = Endo

**Figure S4:**
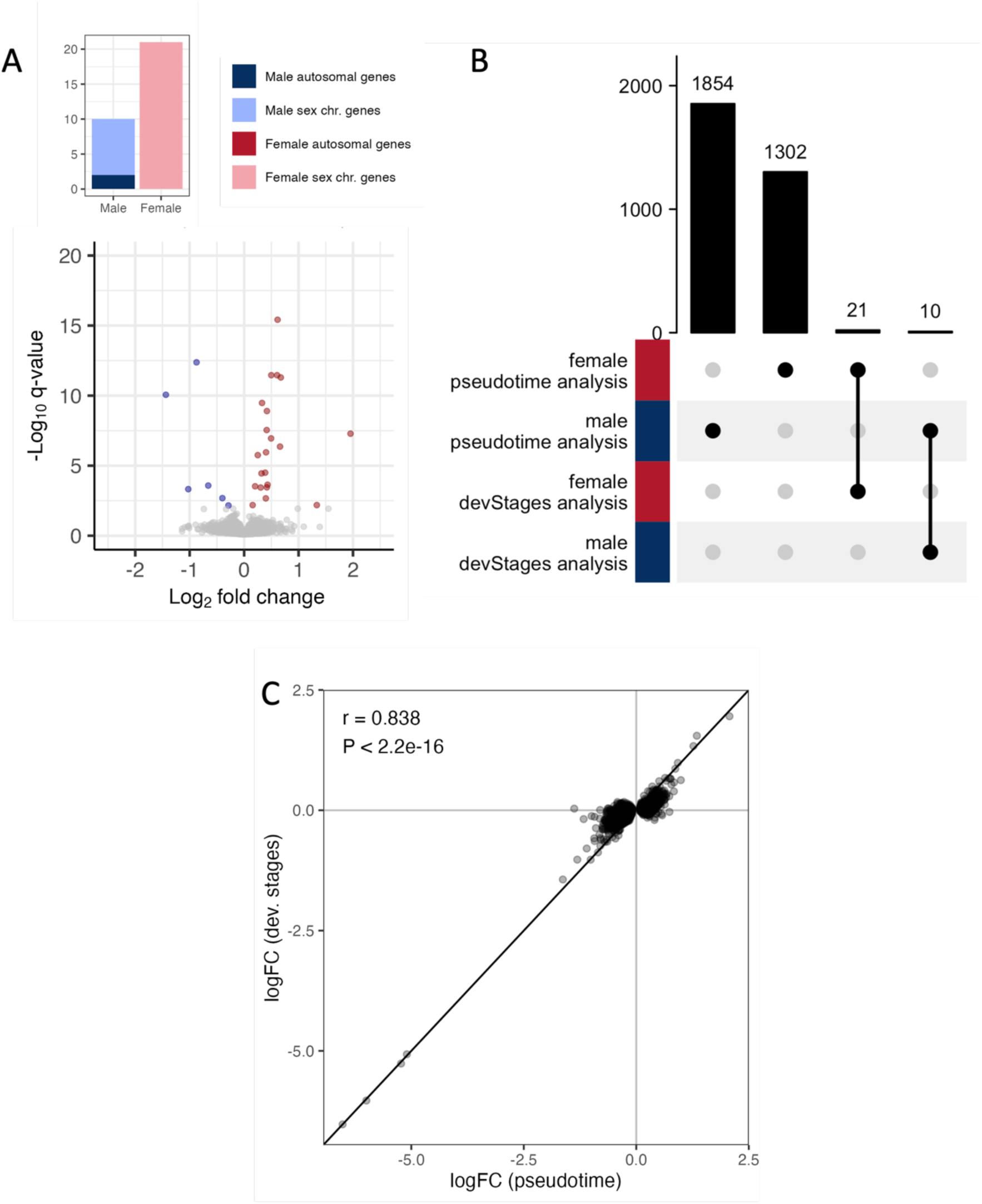
Sex-DE analysis in prenatal dataset with categorical developmental stages as a covariate. (A) Barplot of the number of sex-DE genes and volcano plot (female-biased genes are shown in red and male-biased genes in blue). (C-D) Comparison of the sex-DE analyses with pseudotime and categorical developmental stages as a covariate. (C) Upset plot of the common significant sex-DE genes. (D) Correlation of the effect size (logFC) between the two analyses for genes significantly differentially expressed in the sex-DE analysis using the pseudotime as a covariate.

**Figure S5:**
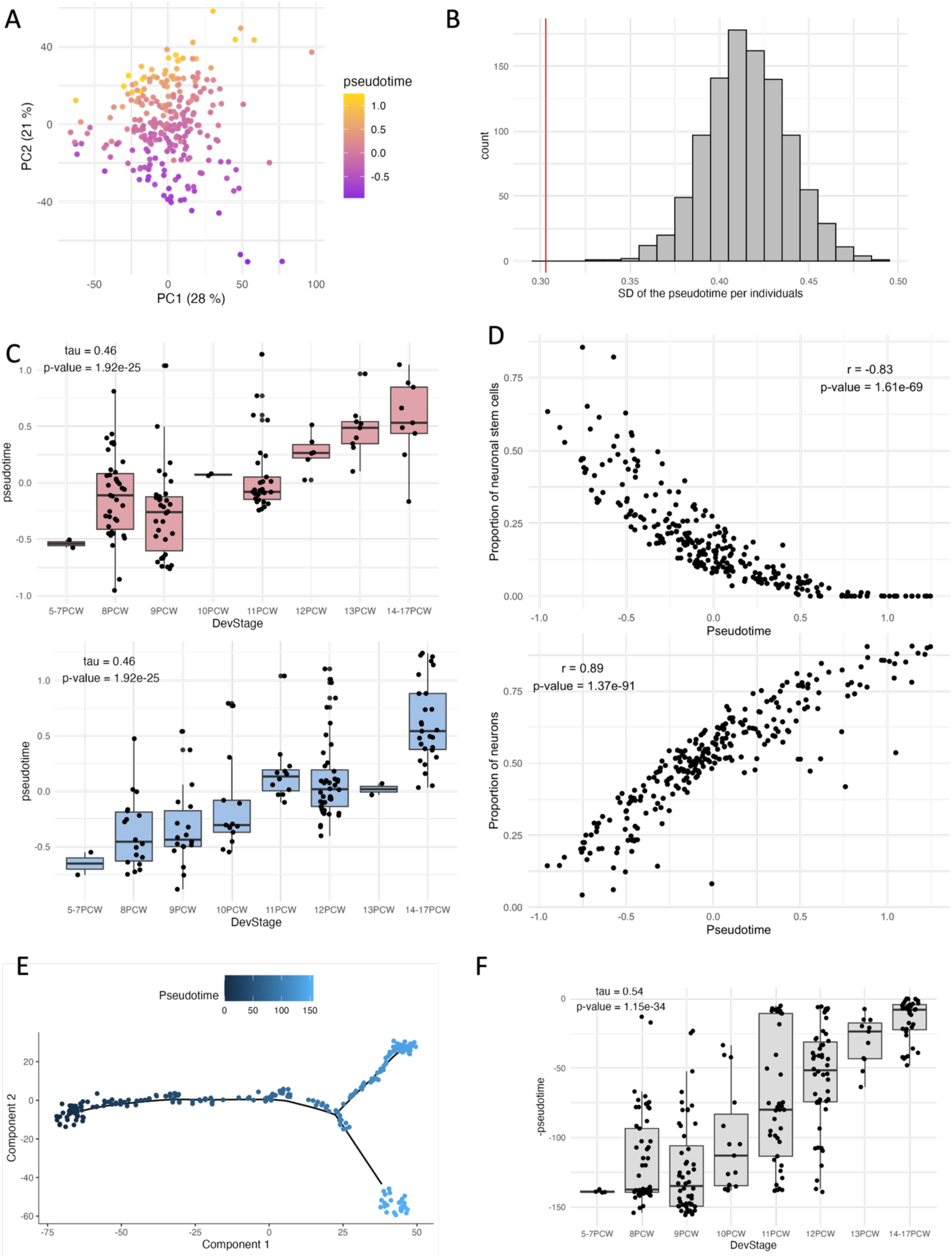
Pseudotime analysis on prenatal data. (A) PCA plot of the prenatal samples colored by pseudotime. (B) Permutation test for the mean standard deviation (SD) of samples from the same individual. (C) Correlation between pseudotime and developmental stages separated by sexes. Top: females, bottom: males. (D) Neuronal stem cells and neurons proportions inferred with CIBERSORTx along pseudotime. (E & F) Pseudotime analysis with monocle. (E) Inferred sample trajectory colored by pseudotime. (F) Correlation between developmental stages and monocle inferred pseudotime.

**Figure S6:**
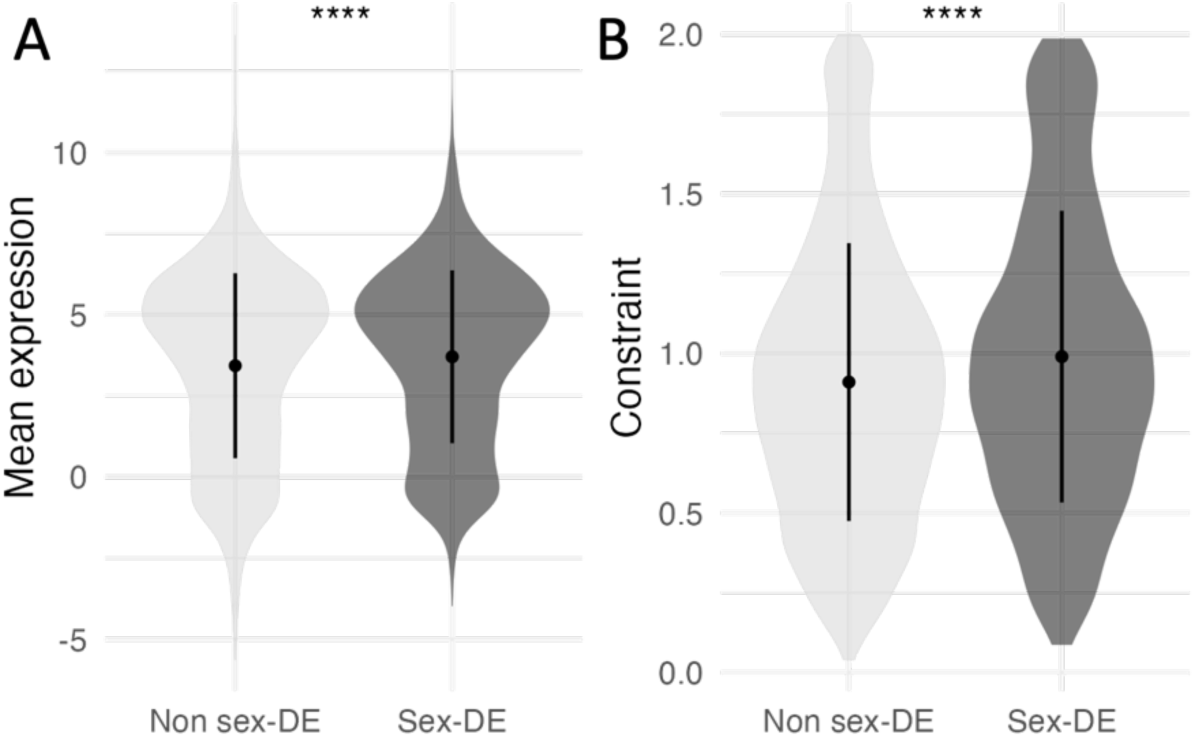
Characteristics for prenatal sex-DE genes (N=3187) compared to non-sex-DE genes (N=14414). (A) Comparison of the average gene expression in prenatal brain between the two groups of genes. (B) Comparison of gene constraint (LOEUF) between the two groups of genes. Smaller LOEUF values indicate higher constraint. Wilcoxon test p-value: **** <= 0.0001.

**Figure S7:**
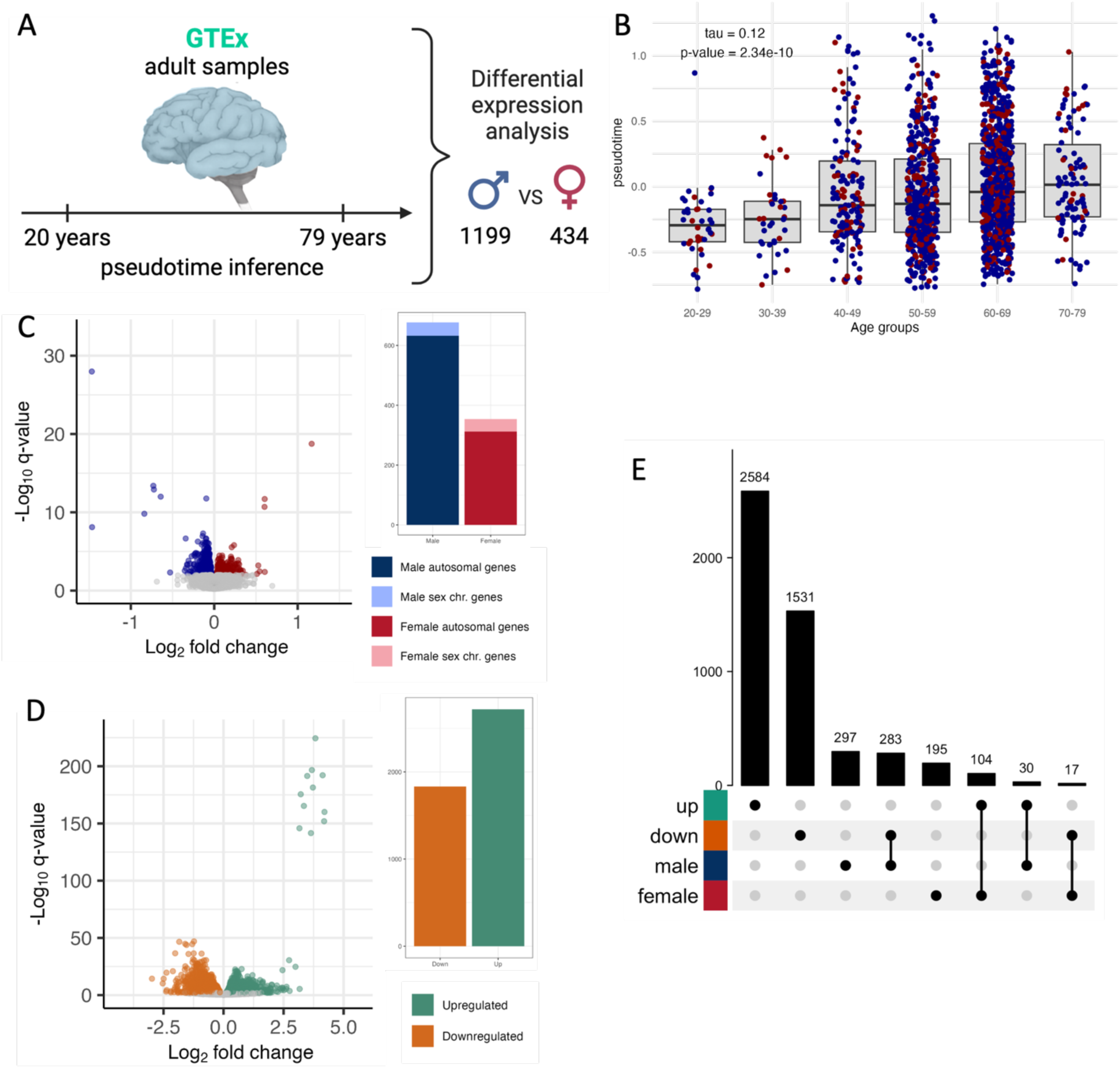
Pseudotime and DE analysis for GTEx forebrain dataset. (A) 1633 adult samples ranging from 20 to 79 years. A pseudotime analysis followed by a differential expression analysis comparing males and females was carried out. (B) Correlation between pseudotime and age groups. (C-D) Barplot of the number of DE genes (q-value < 0.1 and |log2FC| > 0.01) and volcanoplot for the two analysis: (C) sex-DE genes and (D) pseudotime-DE genes. On the sex-DE volcanoplot only autosomal genes are shown. Only significant genes are colored in the volcanoplots. (E) Intersection of the two DE analysis as an upset plot taking into account the direction of effect. The upper barplot shows the size of the intersections.

**Figure S8:**
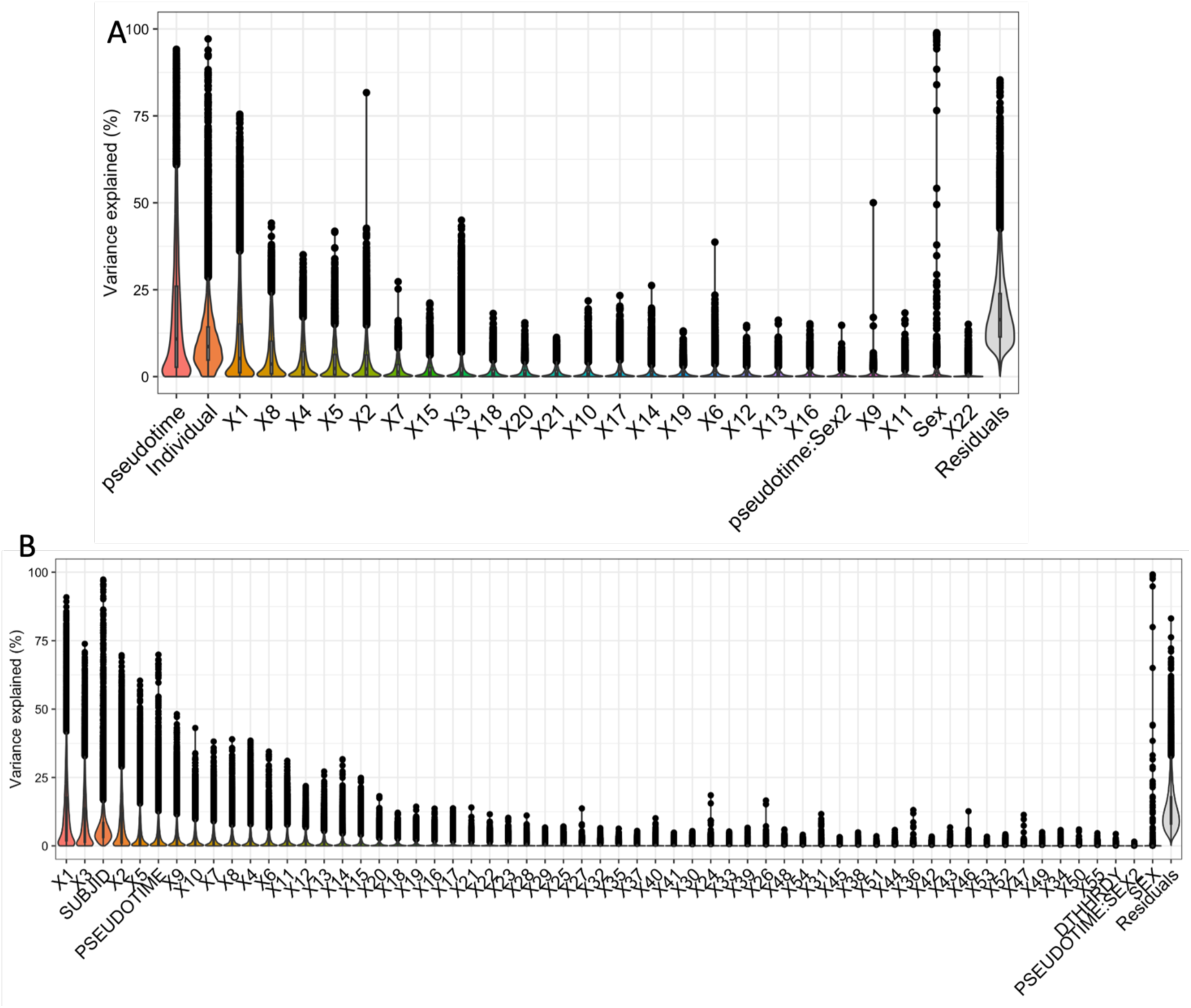
Variance explained of each variable in the model used in the DE analysis for (A) the prenatal dataset and (B) the adult dataset.

**Figure S9:**
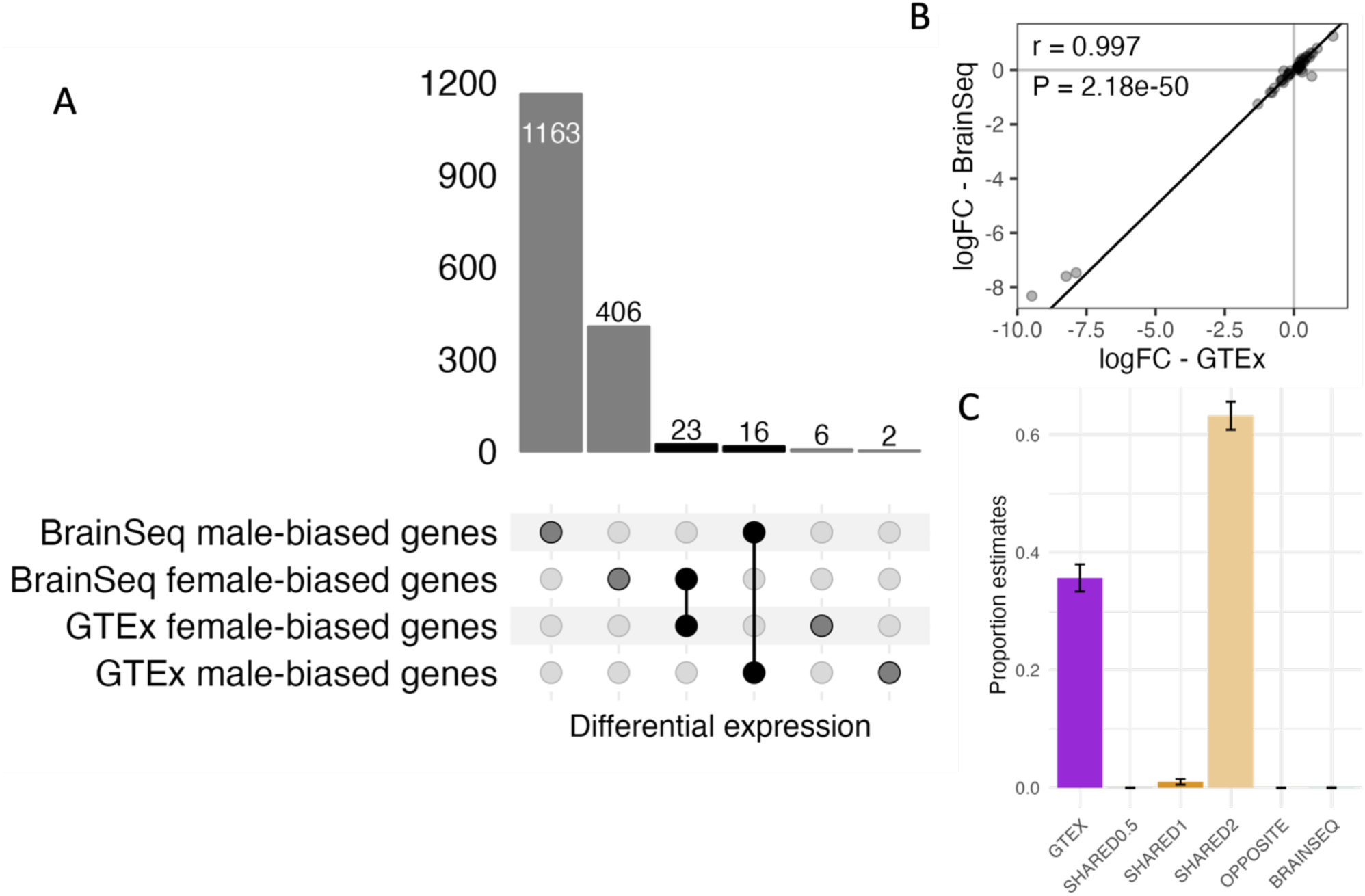
Comparison of sex-DE genes in the same tissue (cortex) for two independent datasets: GTEx prefrontal cortex (N=209) and BrainSeq phase1 DLPFC (N=189). (A) Upset plot showing the overlap of the two datasets accounting for the direction of effect. (B) Correlation of the logFC of the GTEx sex-DE genes between the two datasets. (C) Proportions estimates and SE of each gene categories in the bayesian analysis: GTEx-specific (GTEX), shared with effect size twice larger in GTEx data (SHARED0.5), shared with same effect size (SHARED1), shared with effect size twice larger in BrainSeq brain (SHARED2), opposite effect size and BrainSeq-specific (BRAINSEQ).

**Figure S10:**
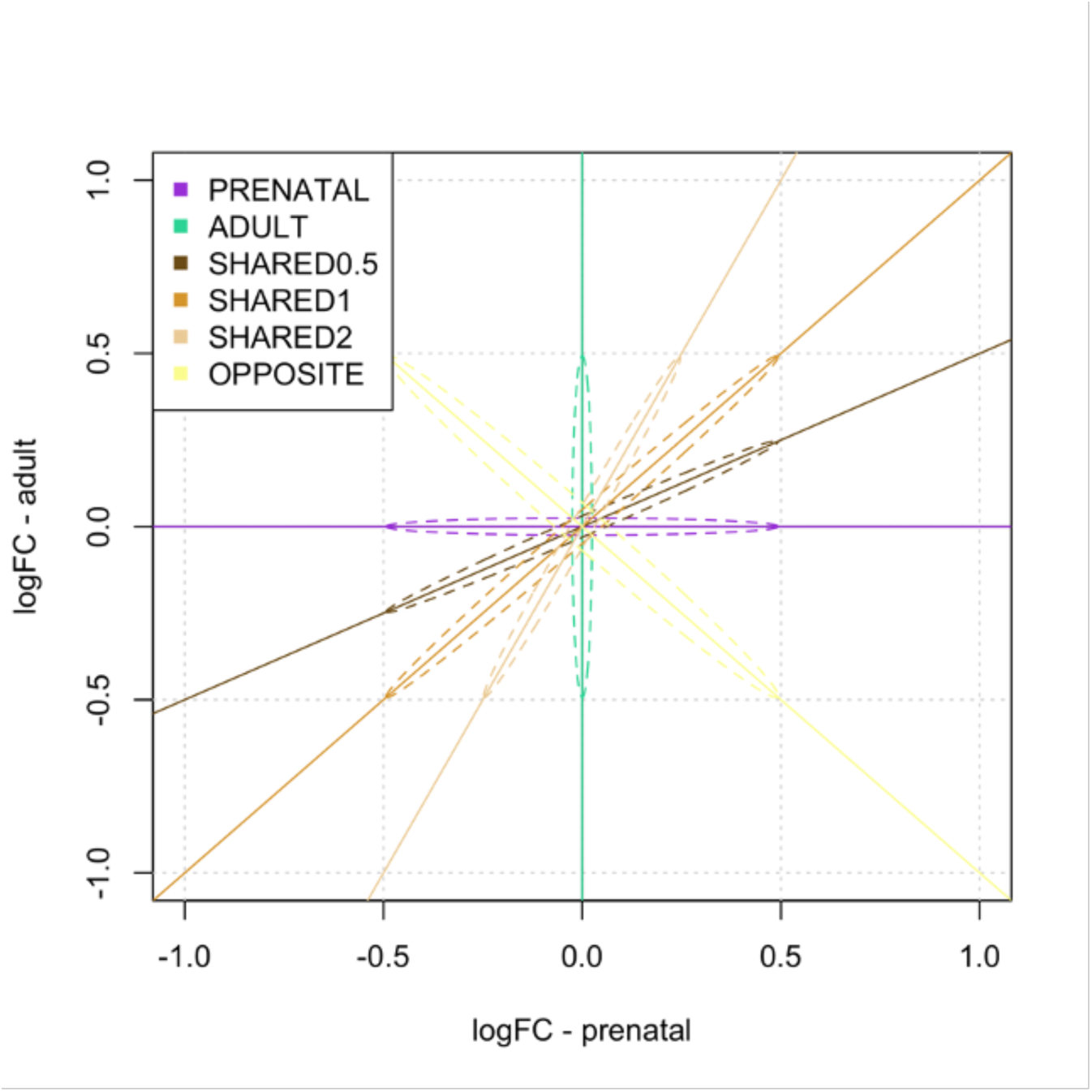
LineModel plot showing the lines of the 6 models used in the Bayesian analysis.

**Figure S11:**
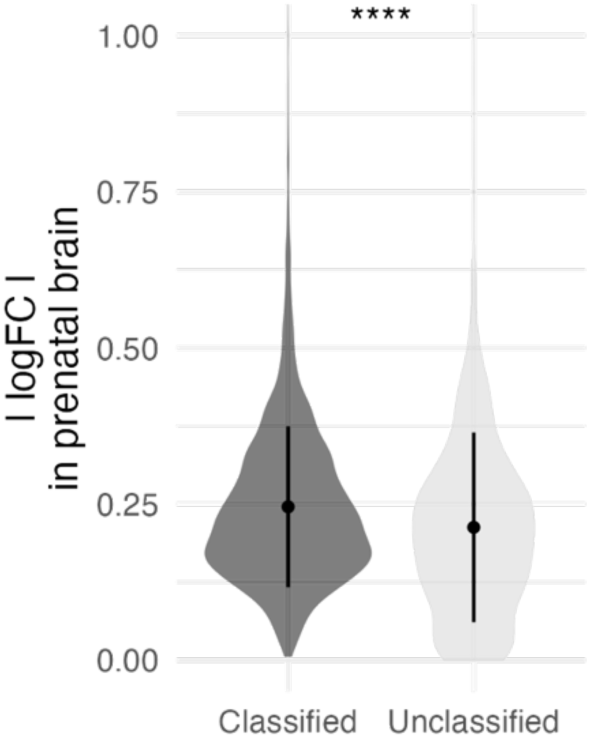
Difference in effect size, |logFC|, in prenatal brain between the sex-DE genes classified in one of the 6 categories and the unclassified ones.

**Figure S12:**
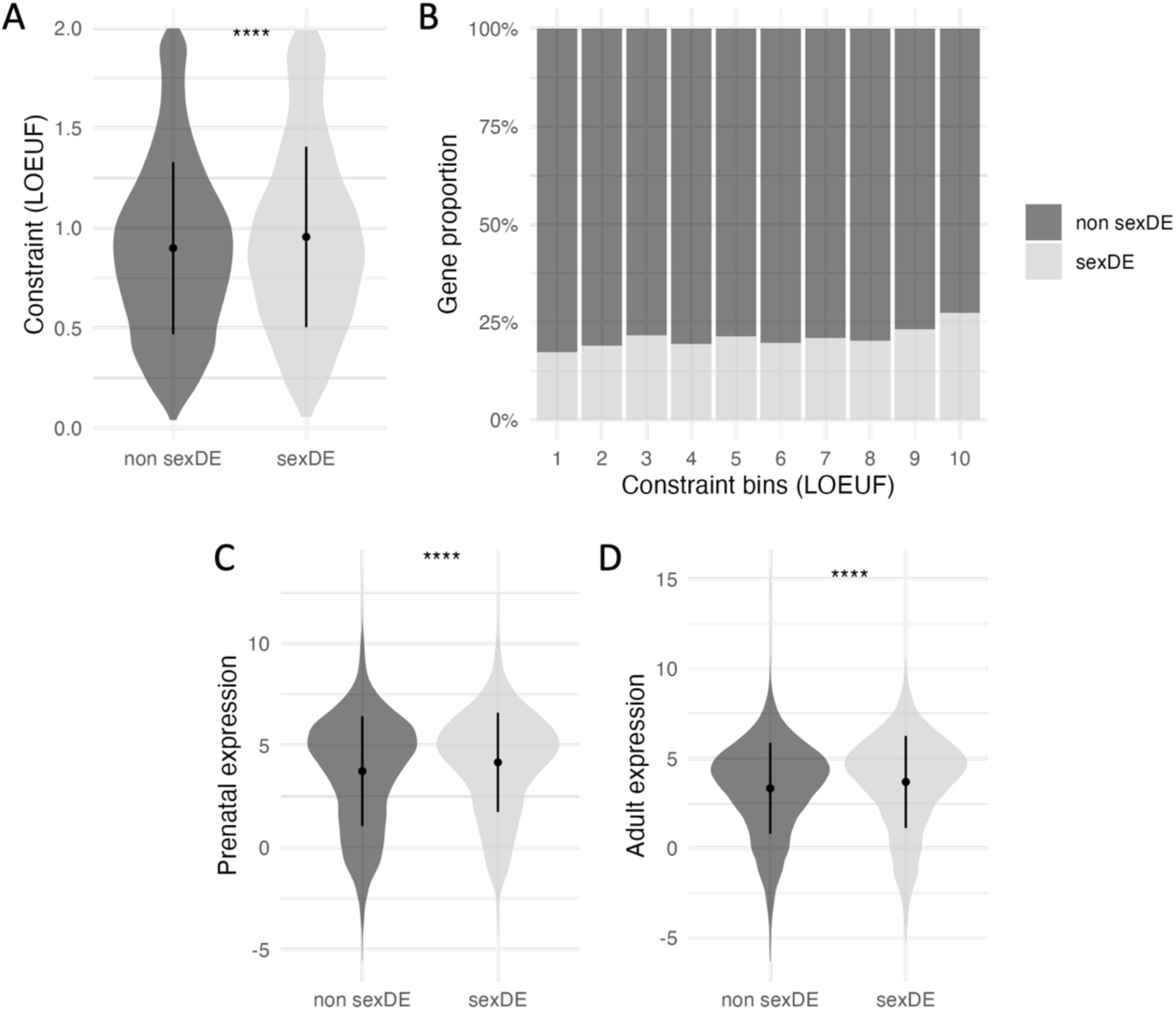
Characteristics of sex-DE genes in either adult or prenatal forebrain (N= 3356) compared to non-sex-DE genes. (A) Comparison of genes constraint (LOEUF) between the two groups of genes. (B) Proportion of sex-DE and non-sex-DE genes in each bin of gene constraint. The bins correspond to the 10 deciles of the LOEUF score. (C-D) Comparison of the gene expression between the two groups of genes in (C) prenatal forebrain and (D) adult forebrain. Wilcoxon test p-value: **** <= 0.0001.

**Figure S13:**
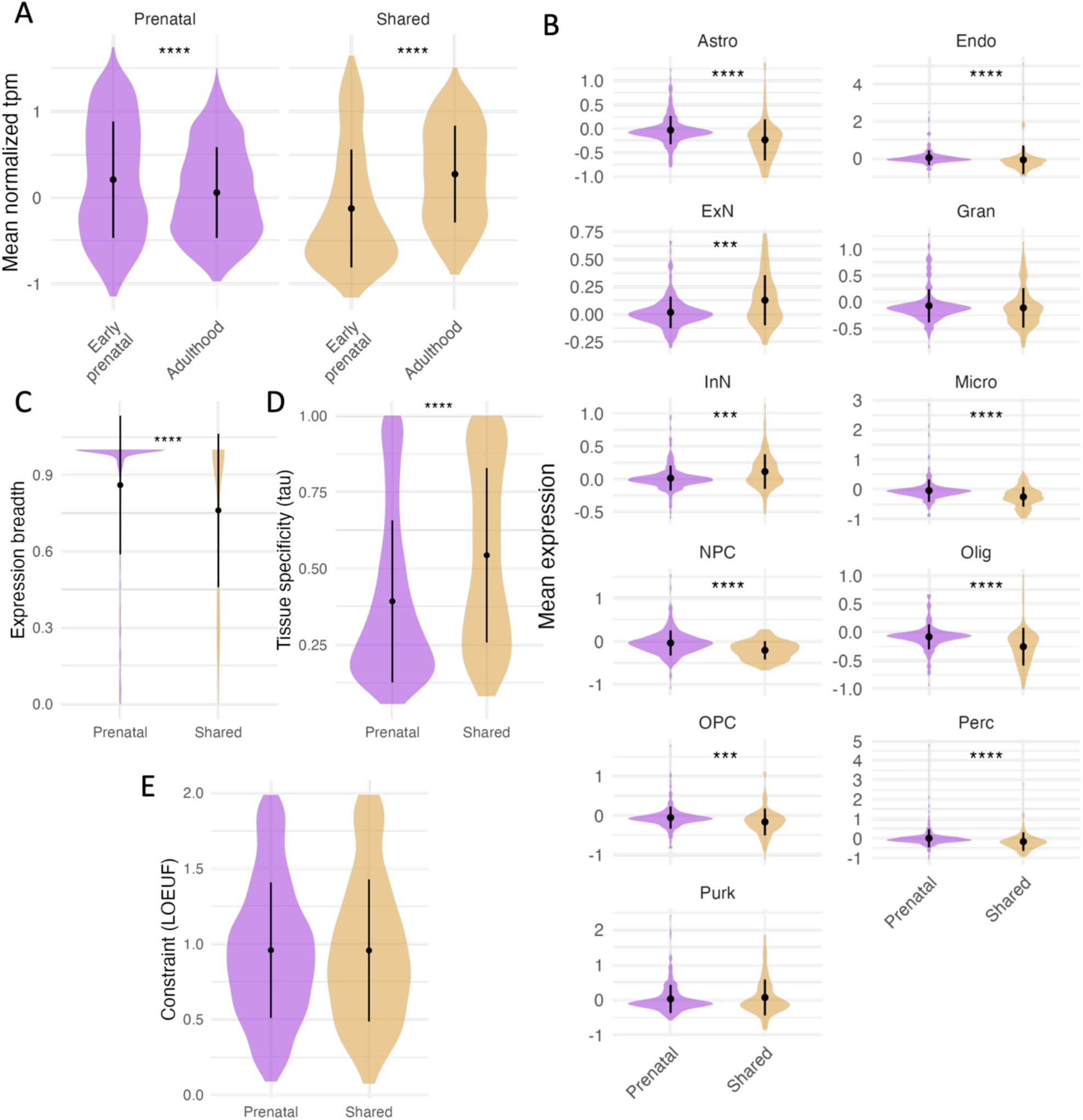
Characteristics of shared and prenatal-specific sex-DE genes between adult and prenatal forebrain. (A) Comparison of the average gene expression by gene category between BrainSpan early prenatal and adulthood samples. (B) Comparison of the average gene expression per gene category in each cell type found in brain single-cell RNA-seq data. (C) Comparison of the expression breadth between the shared and prenatal-specific sex-DE genes. The expression breadth of a gene is defined as the percentage of GTEx tissues in which a gene is expressed (TPM >= 1). (D) Comparison of the tissue specificity (tau value) between the shared and prenatal-specific sex-DE genes. (E) GO term significantly enriched in male-biased prenatal specific and shared genes and in female-biased prenatal specific genes. No significant GO term were found in female-biased shared genes. (F) Percentage of GTEx tissue pairs sharing sex-DE effect (measured with linemodel bayesian framework) with same direction of effect for shared and prenatal-specific genes. (G) Percentage of GTEx tissues showing sex-DE effect (with q-value < 0.01) for shared and prenatal-specific genes. Wilcoxon test adjusted p-value: **** <= 0.0001, *** <= 0.001

**Figure S14:**
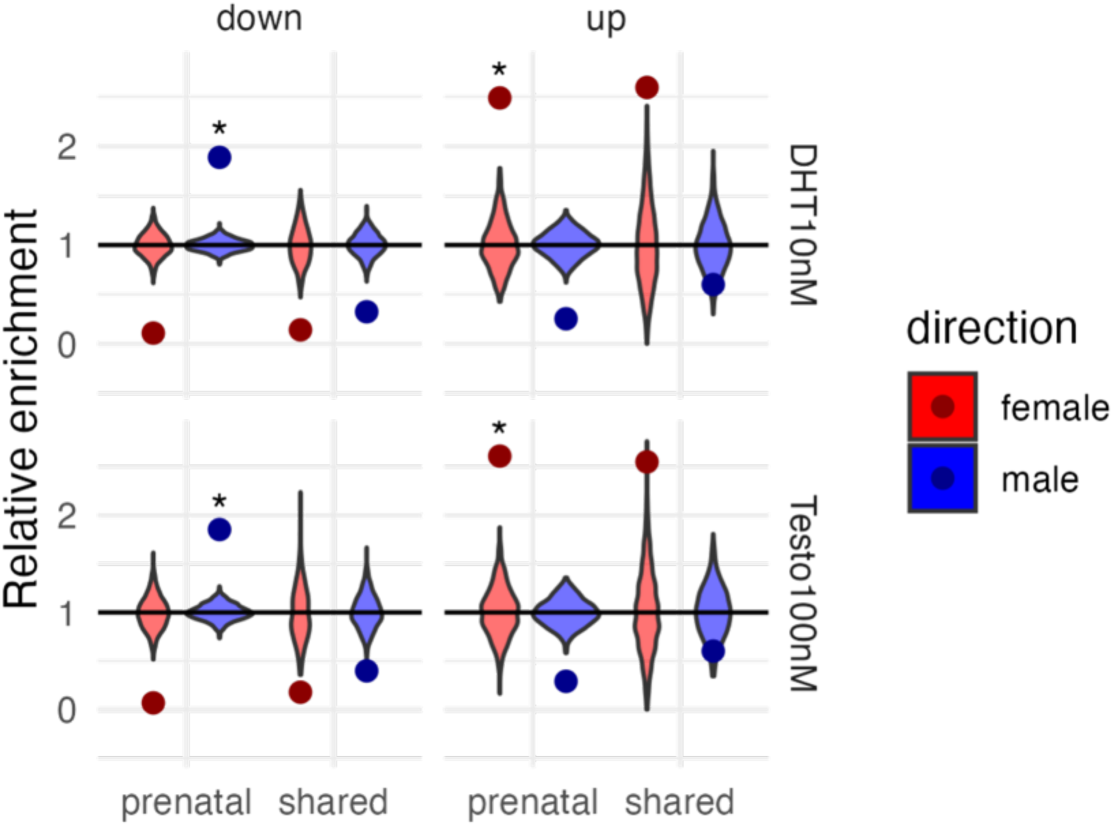
Enrichment analysis of shared and prenatal-specific sex-DE genes with genes regulated by androgen treatments: testosterone (Testo 100nM) and dihydrotestosterone (DHT 10nM). The enrichment of the list of genes of interest (dot) is compared to the enrichment of 1000 random gene lists (violinplot). * denotes the significant enrichments.

**Figure S15:**
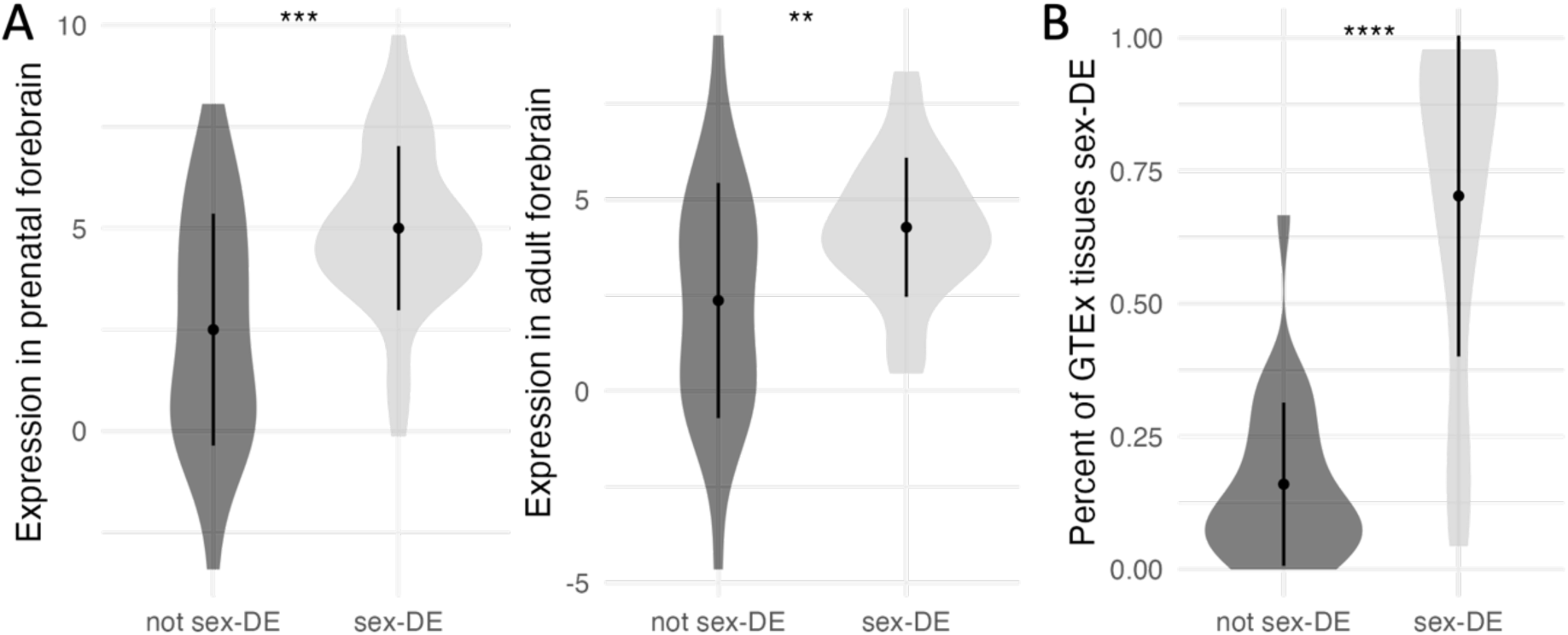
Characteristics of established X-chromosome escape genes found sex-DE in at least one of the two life-stage (N=42) or in neither of them (N=25). (A) Comparison of the average expression of the two groups in either prenatal forebrain (left) or adult forebrain (right). (B) Percentage of GTEx tissues showing sex-DE effect (with q-value < 0.01) for each group of genes. Wilcoxon test p-value: **** <= 0.0001, *** <= 0.001, ** <= 0.01.

**Figure S16:**
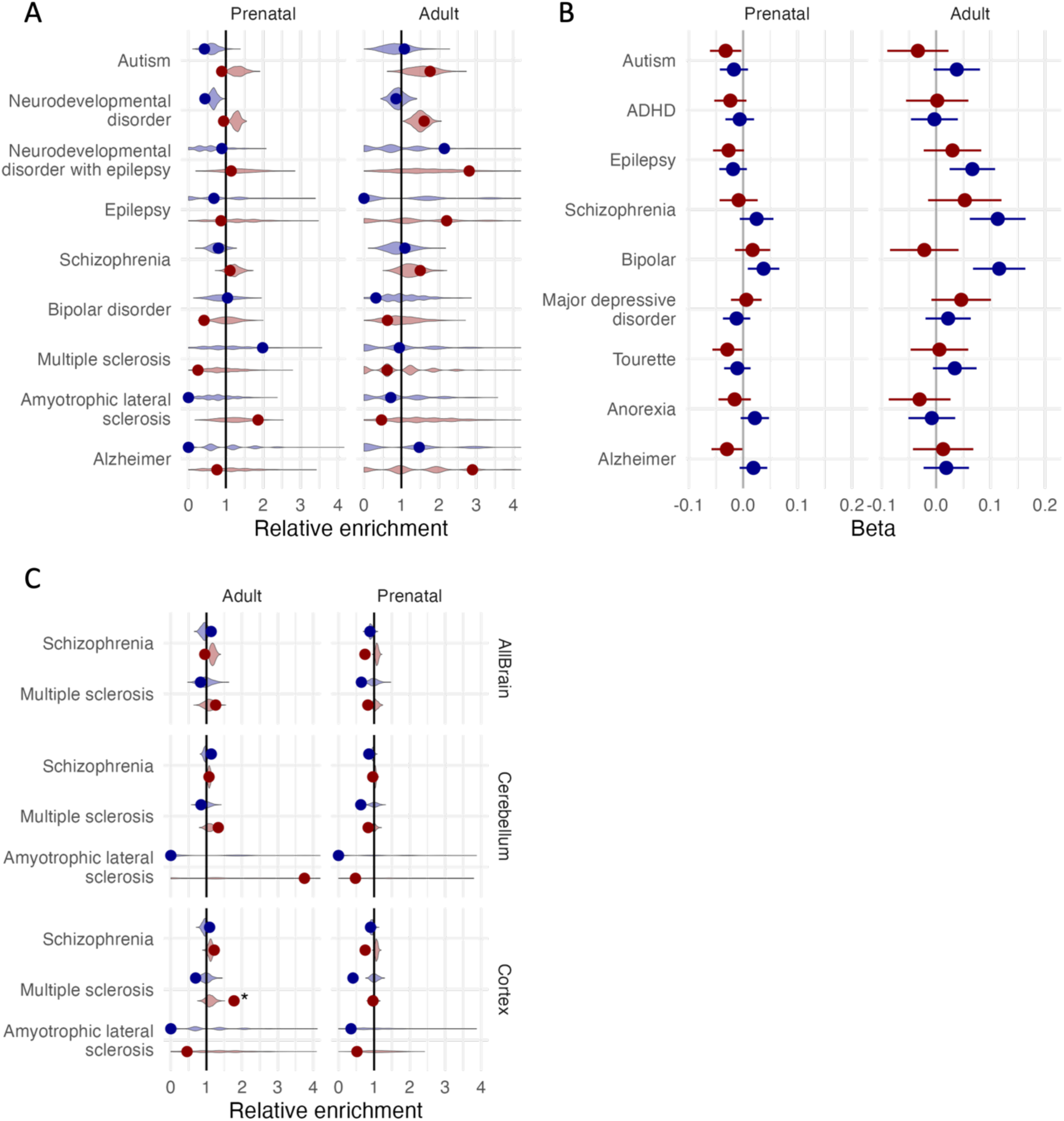
Enrichment analyses for the prenatal and adult sex-DE genes in several neurodevelopmental and neurological diseases. (A) Enrichment analyses with hypergeometric test of gene lists from exome-sequencing studies. (B) Enrichment analyses with the MAGMA method of gene lists from GWAS. The error bars depict the SE of the beta values. (C) Enrichment analyses with hypergeometric test of gene lists from co-regulation networks from the MetaBrain resource. The violinplots on the (A) and (C) plots represent the distribution of the relative enrichment for the 1000 random gene lists and the point the enrichment for the true gene list. *: gene lists passing the significance threshold as defined in Methods.

**Figure S17:**
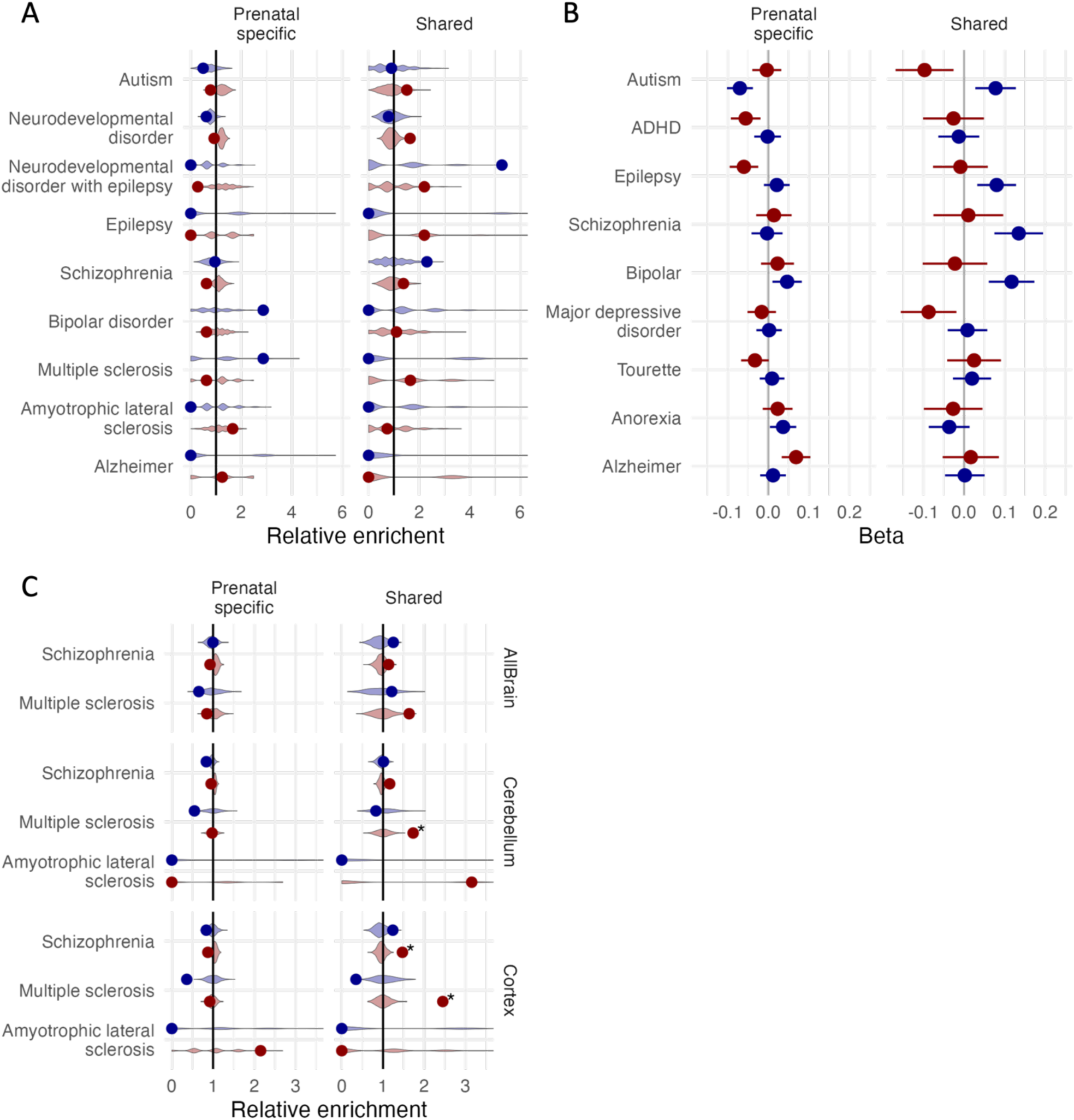
Enrichment analyses for the shared and prenatal-specific sex-DE genes in several neurodevelopmental and neurological diseases. (A) Enrichment analyses with hypergeometric test of gene lists from exome-sequencing studies. (B) Enrichment analyses with the MAGMA method of gene lists from GWAS. The error bars depict the SE of the beta values. (C) Enrichment analyses with hypergeometric test of gene lists from co-regulation networks from the MetaBrain resource. The violinplots on the (A) and (C) plots represent the distribution of the relative enrichment for the 1000 random gene lists and the point the enrichment for the true gene list. *: gene lists passing the significance threshold as defined in Methods.

**Figure S18:**
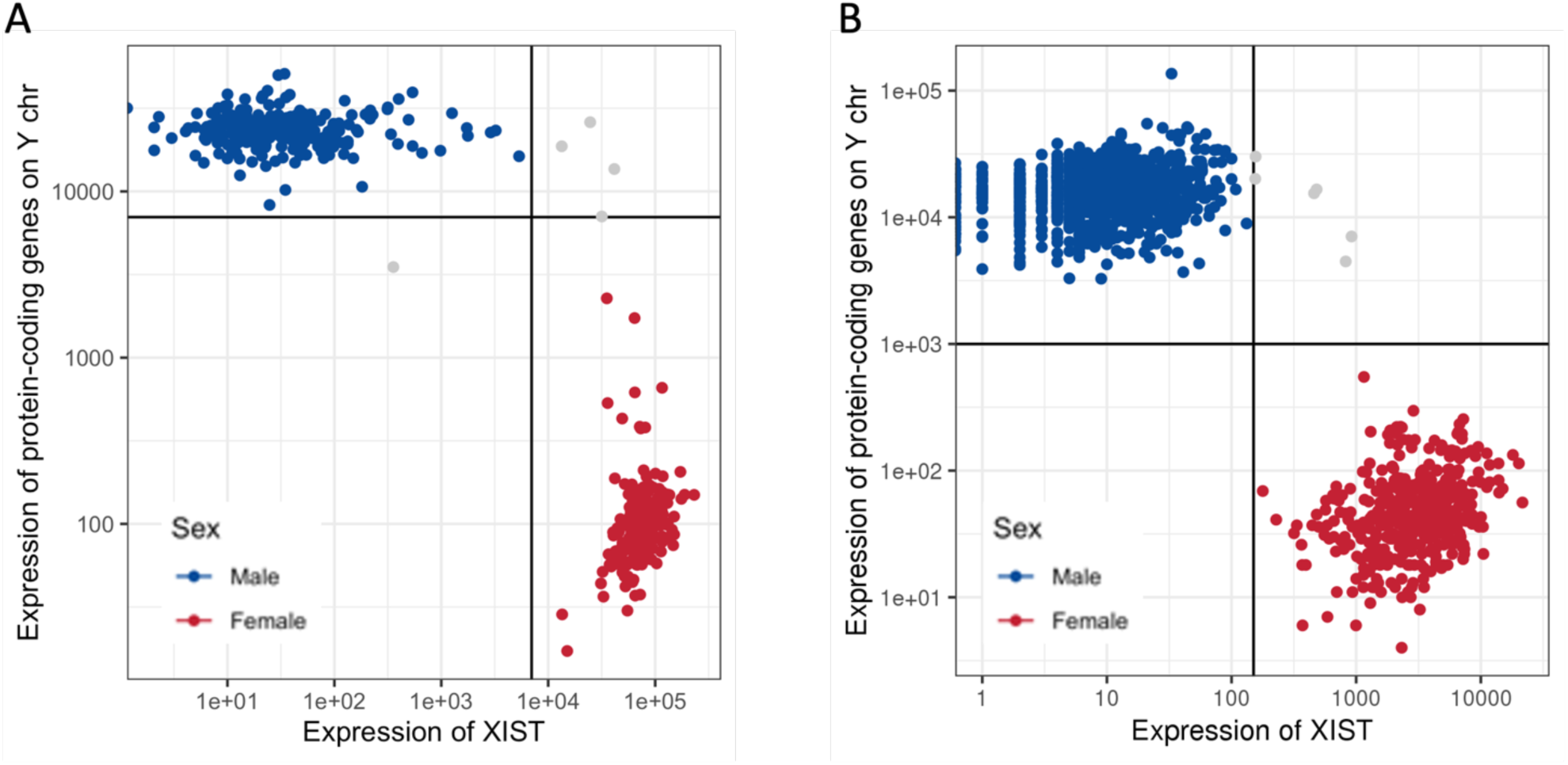
Sex labelling for each sample in (A) the prenatal and (B) the adult datasets using Y chromosome protein coding genes and *XIST* expression.

## Methods

### Data

Three different public RNA-seq datasets were used in this study: Human Developmental Biology Resource (HDBR)^34^, Genotype-Tissue Expression project (GTEx)^35^ and BrainSeq^75^. The number of samples per dataset, per tissue and age group (pre- or postnatal) are detailed in **Table S1**.

#### HDBR dataset

The HDBR dataset contains 542 prenatal RNA-seq samples from 187 distinct individuals. For our analysis, we selected only the samples coming from the forebrain (specific region of the forebrain, forebrain fragment, or whole forebrain). We defined the sex of the individuals by checking the number of reads mapped to genes known to be expressed in one sex only: the protein-coding genes of the Y chromosome (only expressed in males) and *XIST* gene (only expressed in females). A cutoff of 7000 was used to separate male and female samples (**Figure S18A**). Meaning that a sample was labeled male if the count on chromosome Y protein-coding genes was greater than 7000 and the count on *XIST* gene lower than 7000. And vice versa, a sample was labeled female if the count on chromosome Y protein-coding genes was lower than 7000 and the count on *XIST* gene greater than 7000. All samples for which we could not assign sex using these criteria were removed from our analysis. We also removed the only sample from 4PCW and one outlier sample for which the counts were different from all other samples (from Pearson’s correlation analysis). The final number of samples used is 266 corresponding to 72 distinct individuals (**Figure S1A**). The developmental stages we will be referring to in this work correspond to the number of weeks post-conception (PCW) assigned by the HDBR consortium to each fetus. The age of the samples spans from 5 to 17 PCW covering the second half of the first trimester of pregnancy and the beginning of the second trimester. The developmental stages of the earliest samples (up to 60 days post conception) were annotated by HDBR using Carnegie stages, and the different Carnegie stages were merged into PCW. We then divided the samples into 8 different developmental stages as follows: 5-7PCW, 8PCW, 9PCW, 10PCW, 11PCW, 12PCW, 13PCW, and 14-17PCW.

#### GTEx dataset

GTEx contains postmortem adult samples from a wide range of tissues. The forebrain RNA-seq samples were used to assess sex-DE in adult. 1633 samples (434 female and 1199 male samples) from 337 distinct individuals aged between 20 and 79 years and spanning 8 different forebrain regions (amygdala, anterior cingulate cortex, caudate, frontal cortex, hippocampus, hypothalamus, nucleus accumbens, and putamen) were used (**Figure S1B**). Again, the sex of the samples was confirmed using the counts data of *XIST* and the protein-coding genes on the Y chromosome (**Figure S18B**). In this data, a threshold of 150 for the count of *XIST* and 1000 for the sum of the counts of the protein-coding genes on the Y chromosome.

#### BrainSeq dataset

The data from the BrainSeq (phase 1) dataset was used to assess the replicability of the sex-DE genes in the adult cortex. For this replication analysis, we selected only the 189 samples (134 male and 55 female) from dorsolateral prefrontal cortex (DLPFC) of control adults.

### Pseudotime analysis

For each sample, the pseudotime was inferred using the 3000 most highly variable genes with the phenopath R package^40^. The expression values were formatted as log_2_(TPM+1) and the following parameters were used: elbo_tol = 1x10^-6^, thin = 20 and design matrix = ∼Sex. This analysis uses sex as a covariate allowing us to detect different trends in pseudotime between males and females. We checked that the function had reached convergence using the `plot_elbo` function. The set of genes used to infer pseudotime was removed from all analyses related to pseudotime (i.e pseudotime-DE analysis).

To compute the Kendall correlation between the inferred pseudotime and the developmental stages, we converted the categorical variable corresponding to the developmental stages to a continuous variable.

### Cell type decomposition analysis

We used CIBERSORTx^38^ to do a cell type decomposition of the prenatal and postnatal forebrain samples. This tool uses single-cell RNA-seq (scRNA-seq) data to infer the transcriptomic signature of each cell type and then estimates the abundance of each cell type in the bulk RNA-seq samples. We used 3 publicly available scRNA-seq datasets present in the STAB database^48^ as the reference. The 3 chosen datasets^76–78^ contained samples from embryonic (5 to 26 PCW) and adult (22 to 63 years) individuals and spanned different regions of the forebrain (**Table S1**). We used the docker container to run CIBERSORTx with the following parameters: `--rmbatchSmode TRUE --perm 100 -- replicates 5 --sampling 0.5 --fraction 1 --k.max 999 --q.value 0.01`.

### Normalization of expression counts

For the HDBR forebrain dataset, transcripts abundance was assessed with Salmon^37^ using Gencode (version 28) reference annotation. Counts for each transcript of a gene were summed to obtain the gene quantification. These counts were then normalized using EdgeR^79^ TMM normalization. Only genes with count per million (CPM) values higher than 1 in more than 10 samples were kept for the differential analysis. Finally, the normalization was done with `voomWithDreamWeights` function from variancePartition R package^41,42^ using sex, pseudotime, individuals, and surrogate variables (svas) as covariates. The svas were computed with sva R package^80^.

For the GTEx dataset, we used the raw counts (using version 26 of Gencode annotation) publicly available on the online portal (https://www.gtexportal.org/home/). We did the same TMM normalization with EdgeR, applied the same filter on CPM values, and carried out the normalization with `voomWithDreamWeights` using sex, pseudotime, cause of death, individuals, and svas as covariates.

### Sex differential expression analysis

We tested for genes differential expression between males and females defined as a binary traits using the linear mixed model for repeated measures (dream) from the variancePartition package^41,42^, adjusted for covariates, i.e., pseudotime, pseudotime/sex interaction, and surrogate variables as fixed effects, and accounting for multiple samples from the same individuals as a random effect.

In the forebrain HDBR analysis, the following model was used:

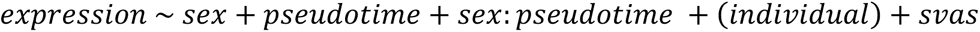

In the forebrain GTEx analysis, the following model was used:

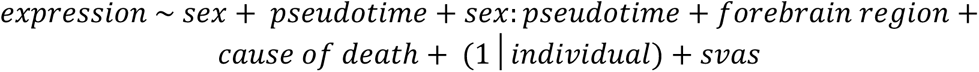

For both datasets, we extracted three lists of DE genes:

- genes differentially expressed between males and females (sex-DE)
- genes differentially expressed along pseudotime (pseudotime-DE)
- genes with a significant interaction between sex and pseudotime (interaction-DE) For all genes in each analysis, the false discovery rate (q-value) was computed using the qvalue R package^81^ to correct for multiple testing. In all these analyses, a gene was considered significantly differentially expressed if it passed the following thresholds: q-value < 0.01.

Sex differencial expression refers to group-level average differences between individuals defined as males or females in our analysis.

### Comparison of sex-DE genes across life stages or datasets

To compare the results of the differential analysis between the two life stages studied, we first look at the overlap of the sex-DE genes (q-value < 0.01). A hypergeometric test was used to assess the significance of the overlap. For the comparison not to be too impacted by the choice of the thresholds and by the power of each dataset, we also computed the replication rate (π_1_) using the qvalue R package^81^ with `lambda = 0.5`.

We also computed a consistency metric for the sex-DE. This metric represents the percentage of genes that vary in the same direction in the sex-DE analysis (the sign of the t-statistic is the same in both analyses). We tested if the consistency was different than what was expected by chance using a binomial test. The consistency of the sex-DE effect size was also assessed with Pearson’s correlation and visualized in a scatter plot.

These analyses were performed on all sex-DE genes and then, we separated the genes into three groups: autosomal genes, genes escaping XCI, and inactive X chromosome genes as defined in^57^.

Similar metrics were computed to compare two dataset of the same tissue and same life-stage: GTEx frontal cortex (BA9) and BrainSeq DLPFC.

### Bayesian method to compare sex-DE across dataset

To compare sharing and specificity of sex-DE genes between prenatal and adult brain, we used the *linemodel* Bayesian method^47^. For all sex-DE genes (prenatal or adult), we estimated probabilities between six possible explanations of the observed data: the effect is present in only one of the dataset but not in both (PRENATAL *slope*=0 and ADULT *slope*=Inf); the effect is shared by the datasets with the same effect size (SHARED1 *slope*=1); the effect is shared with a higher effect size in prenatal brain (SHARED0.5 *slope*=0.5); the effect is shared with a higher effect size in adult brain (SHARED2 *slope*=2); or the datasets have opposite effect sizes (OPPOSITE *slope*=-1) (**Figure S10**). The prior variance of effect size (scale) was set to 0.1971916 for all models which correspond to the 95th percentile of the logFC in the prenatal dataset divided by 2. We set the correlation parameter (*cor*), which corresponds to the allowed deviation from the expectation to 0.995 for all models except for OPPOSITE for which *cor*=0.990. Lastly, the correlation (*r.lkhood*) between the estimators of the effect sizes between the two dataset was set to 0 as the datasets are independent (i.e. they don’t contain any overlapping individuals). We used the *line.models.with.proportion* function that gives for each gene a posterior probability on each model as well as the overall proportions of effects coming from each model with 2000 iterations and a burn-in period of 200 iterations.

For each gene, we then classified them into 4 categories using threshold on the posterior probability: shared effect (SHARED0.5 + SHARED1 + SHARED2 > 0.8), prenatal-specific effect (PRENATAL > 0.8), adult-specific effect (ADULT probability > 0.8) and opposite effect (OPPOSITE > 0.8).

This Bayesian approach with the same parameters was also used to compare sex-DE results of all GTEx tissues against each other as well as the GTEx forebrain dataset against the BrainSeq dataset. For the comparison of GTEx tissues, to account for the fact that some samples from different tissues could come from the same individual, we computed a correlation between the pairs of tissues. For each pair of tissue, the *r.lkhood* parameter was computed by taking the Pearson correlation between the logFC of all genes that are not strongly sex-DE (pvalue > 0.10) in both tissues.

### Analysis of the sex-DE genes properties

We looked at several properties of the sex-DE genes: their level of expression in both bulk and single-cell data, their level of constraint, their breadth of expression, tissue specificity and consistency in sex-DE across adult human tissue (GTEx).

For the comparison of genes level of expression, we used expression from bulk RNA-seq data in adult and prenatal brain samples from BrainSpan (www.brainspan.org). We also used the genes expression from the scRNA-seq datasets from the STAB database^48^ previously describe, allowing to look at expression in different cell types. For the comparison of genes constraint, we used the LOEUF score describe in^60^ downloaded from Gnomad (v4.1) browser. In addition, we used several metrics derived from the full GTEx (v8) dataset containing gene expression for 52 human adult tissues. We compared the breadth of expression, i.e. in how many tissues a gene is expressed in, the tissue specificity (tau value) computed by Palmer and colleagues^82^, and the consistency of sex-DE across tissues, i.e. in how many tissues a gene is sex-DE with the same direction of effect (male- or female-biased).

### Enrichment analyses

For all enrichment analyses requiring a list of significant genes, we tested each DE analyses separating the significant DE genes (q-value < 0.01) in two lists corresponding to the direction of effect (either positive or negative logFC). When analyzing the sex-DE genes, the genes used to assign sex to the samples were removed. For the analyses on the Bayesian results, we used the classification of the genes created using the posterior probabilities.

#### GO term over-representation analysis

The over-representation analysis was performed with compareCluster function from the clusterProfiler R package for each GO category separately (Biological process, Molecular Function and Cellular Component)^83^.

#### Enrichment for transcription factor binding sites in promoters

We used the UniBind enrichment tool^51^ to predict if promoters of DE genes were enriched in transcription factor binding sites (TFBS). Promoters were defined as the region spanning 2kb upstream of the gene transcription start site similarly to^16^. We used the subcommand oneSetBg of the UniBind_enrich script with the promoters of all genes analyzed in the differential analysis as background and the robust human TFBS sets computed by Unibind as a database.

#### Enrichment for androgen responsive genes

We performed an enrichment analysis of the androgen responsive genes from Quartier and colleagues^55^ in the prenatal-specific and shared sex-DE genes defined with the Bayesian model. For these analyses, we separated the genes lists by direction of effect (female- or male-biased). We tested the enrichment using a hypergeometric test and then, we carried a permutation analysis (1000 permutations) to determine if the enrichment or depletion seen is more than expected by chance. An enrichment was considered significant if the hypergeometric p-value was lower than 0.0005 and the permutation p-value was lower than 0.05.

#### MAGMA analysis

The MAGMA^84^ gene-set analysis was carried on the significant sex-DE genes (q-value < 0.01) separating the male- and female-biased genes both for prenatal and adult forebrain. The same analysis was also carried out on shared sex-DE genes, and prenatal-specific sex-DE genes (again separating the male- and female-biased genes). Nine disease traits were analyzed: ADHD^85^, Alzheimer^86^, Anorexia^87^, ASD^88^, Bipolar disorder^89^, Epilepsy^90^, Major depressive disorder^91^, Schizophrenia^92^ and Tourette syndrome^93^. The MAGMA analysis test whether the genes from the different gene sets are more strongly associated with the phenotypes of interest compared to other genes. We mapped the HapMap3 single nucleotide polymorphisms to protein-coding and lincRNA genes within 50 kb upstream and downstream, using the 1,000 Genomes Phase 3 European dataset as the reference for variant locations and Gencode annotation (version 26) as the reference for gene locations. For all other steps, we followed the standard MAGMA pipeline. We filtered significant enrichment using the p-value threshold of 6.9x10^-04^ (0.05/9x8) to account for multiple testing.

#### Hypergeometric enrichment analysis

We performed an enrichment analysis on sets of genes linked to neurological or neurodevelopmental disorders through large exome-sequencing studies. Meaning that most genes in these lists were genes where loss-of-function variants were found to be associated with the pathology. The four gene sets are the following: SCZ (Schizophrenia exome meta-analysis consortium^94^, 244 genes), epilepsy^95^ (15 genes), neurodevelopmental disorders with epilepsy^96^ (33 genes), neurodevelopmental disorder^97^ (662 genes), ASD^97^ (183 genes), bipolar disorder (Bipolar Exome sequencing project, https://bipex.broadinstitute.org/, 85 genes) and multiple sclerosis (MS)^98–100^ (28 genes). We also choose two control gene sets from neurological disorders with older age of onset: Alzheimer^101^ (20 genes) and amyotrophic lateral sclerosis (ALS)^102^ (36 genes). In addition, we did an enrichment analysis on the gene lists from the MetaBrain co-expression network^63^ for which at least 5 genes passed the 5% FDR threshold from the downstreamer analysis. We have gene lists for the 3 following diseases: SCZ, MS and ALS.

For each gene set, we computed the relative enrichment in prenatal and adult sex-DE genes as well as on the prenatal-specific and shared sex-DE genes defined with the Bayesian model. For all these analyses, we separated the genes lists by direction of effect (female- or male-biased). We tested the enrichment using a hypergeometric test. As the sex-DE genes are overall more highly expressed than non-sex-DE ones (**Figure S6; Figure S12C-D**), we accounted for this confounder (i.e. level of expression) in our enrichment analysis through matching our background set of genes to genes with similar level of expression in the brain. Finally, we carried a permutation analysis to determine if the enrichment or depletion seen is more than expected by chance.

We filtered significant enrichment and depletion using the hypergeometric test p-value with a threshold of 6.9x10^-04^ (0.05/9x8) and 7.8x10^-04^ (0.05/8x8) for the enrichment of exome-sequencing studies gene lists and the enrichment of the MetaBrain co-expression network gene lists, respectively, to account for multiple testing. In both cases, we also applied the threshold of 5% on the permutation p-value.

