## Supplemental Document 1 for "Early establishment and life course stability of sex biases in the human brain transcriptome"

### Principal component analysis

The principal component analysis (PCA) was carried out using ade4 R package[1] on a total of 4137 samples from pre- and postnatal origin, as well as different tissue origins (**Table S1**). The samples came from the 3 datasets used in the main text (HDBR, GTEx, and BrainSeq) and the EvoDevo mammalian organs dataset[2]. This last dataset comprises 254 samples spanning different tissues (brain, liver, heart, and kidney) and different life stages (from prenatal to adulthood). This dataset was added in the PCA analysis to ensure we were not seeing an effect of the life stage confounded by the dataset of origin. Indeed, EvoDevo is the only dataset containing both prenatal and postnatal samples for each tissue.

The raw counts were first normalized using the variance stabilizing transformation (vst) function from DESeq2[3]. Then, we removed the effect of the study of origin using the batch correction method removeBatchEffect from limma[4].

In the principal component analysis with samples from all tissues and life stages (**Figure S2A**), we observe a clear separation between the different tissues without a clear distinction between pre- and postnatal samples. While the two subregions of the brain cluster together in this analysis (**Figure S2A**), if we analyze only the brain samples, we see that they have a distinct transcriptomic profile in adults (**Figure S2B**). Finally, focusing only on the forebrain samples, we observe a clear distinction in the transcriptomic profile of the forebrain between the pre- and postnatal life stages (**Figure S2C**).

### Pseudotime analysis with monocle

We used Monocle2 following the "Constructing single cell trajectory" tutorial from this page: http://cole-trapnell-lab.github.io/monocle-release/docs/#trajectory-step-1-choose-genes-that-define-a-cell-s-progress.

First, we chose the genes to use to construct the trajectory. We selected the top 1000 genes DE with the developmental stages. Next, we reduced the dimensionality of the data controlling for the forebrain region as we didn't want it to contribute to the trajectory. Then, we ordered the samples along the created trajectory and visualized the sample colored by pseudotime in a 2-dimensional space (**Figure S5E**). Similarly to what was done with the pseudotime inferred using phenopath, the Monocle2 inferred pseudotime variable correlated well with the developmental stages specified in the metadata (Kendall’s tau=0.54; **Figure S5F**), suggesting again that this time of analysis accurately captures the trajectory of early brain development.

### Pseudotime inference unbiased by sex covariate

Given the systematic sex differences in the developmental trajectory of the brain, we wanted to ensure that the inclusion of the sex covariate did not introduce bias into the pseudotime inference using the phenopath method. To test this, we assigned a random binary variable (1 or 2) to each individual. We then repeated the entire analysis (following the same steps and using the same parameters) that we had performed with the sex variable, but this time using this random variable instead. This included both the pseudotime inference and the differential expression (DE) analysis. We did these analyses 50 times.

We compared the significant pseudotime-DE genes from our main analysis (which used the sex variable) with the new pseudotime-DE results. Remarkably, on average, 92.02% of the significant pseudotime-DE overlapped with those from the original analysis, and the correlation of the effect sizes (logFC) was very high (mean Pearson’s r=0.95). These results indicate that the pseudotime inference is robust, regardless of the covariate used.

### Comparison of prenatal and adult sex-DE genes

Several complementary methods were used to compare sex-DE genes between early development and adulthood.

Focusing on genes with significant sex-DE in either of the two data sets (3187 in the prenatal brain, 1033 in adult brain), we observed a more than two-fold enrichment with concordant effect directions between the two analyses (hypergeometric test p-value=$4.84\times10^{-53}$ for male, 227 shared DE genes; and $7.56\times10^{-06}$ for female, 54 shared DE genes; **Figure 2A**; **Table S5**). Further indicating sharing of the sex-DE patterns, almost half of the prenatal sex-DE genes were estimated to display non-null p-values in the adult brain (replication rate, $\pi_1$=0.46; see Methods), these genes also displayed more consistency in effect directions in the adult brain than expected by chance (72.2% consistency, permutation test p-value=0.001, 1000 permutations) (**Table S5**), and the correlation of the effect sizes (logFC) indicated a good concordance in the magnitude of sex-DE (Pearson's r=0.60, p=4.77x10$^{-261}$, for prenatal sex-DE genes; r=0.35, p=1.04x10$^{-74}$, for autosomal prenatal sex-DE genes; **Figure 2B**; **Table S5**). Using the adult sex-DE genes as the input, these same metrics pointed to an even greater similarity of sex effects between the two life stages (78.3% consistency, Pearson's r=0.84, $\pi_1$=0.68; **Table S5**). The similarity of sex-DE, was, however, lower than in two independent adult cortex data, i.e., datasets of the same tissue and age range (95.7% consistency, Pearson's r=0.99, $\pi_1$=0.87) (**Figure S9A-B**; **Table S5**), suggesting also a degree of distinct sex effects between the two life stages.

Then, we applied a Bayesian model comparison framework[5,6] to estimate the overall proportions of genes consistent with shared, opposite, prenatal-specific, or adult-specific sex-DE patterns and to additionally classify individual genes into different categories of sex-DE including life stage-specific effects and three types of shared effects (equal effect size, SHARED1; effect size twice larger in the prenatal brain, SHARED0.5;  effect size twice larger in the adult brain, SHARED2) (see Methods; **Figure S10**).

Most of sex-DE effects were either prenatal-specific (64.6%) or shared between both life stages (30.6%), leaving only a small fraction of the effects compatible with opposite sex-DE effects (4.2%) and adult-specific sex-DE (0.7%). The latter finding is striking considering that, based on the stringent q-value cutoff of 0.01, 20.2% of the input genes were significantly DE only in the adult brain. Upon closer examination, the 679 genes displaying sex-DE only in the adult data were, while non-significant, enriched in small p-values in the prenatal DE results ($\pi_1$=0.56) but showed increased variance (comparison of logFC standard error for adult only sex-DE genes vs common sex-DE genes, Wilcoxon test p-value=6.52x10$^{-06}$) compared to genes that are sex-DE in both data sets, partly explaining why these genes fail to reach significance for DE but are classified as shared genes by the model.
